# Laminar Architecture of a Decision Circuit in Orbitofrontal Cortex

**DOI:** 10.1101/2025.03.03.641234

**Authors:** Alessandro Livi, Manning Zhang, Mary Carter, Heide Schoknecht, Andreas Burkhalter, Timothy E. Holy, Camillo Padoa-Schioppa

## Abstract

During economic choice, different neurons in orbitofrontal cortex (OFC) encode individual offer values, the binary choice outcome, and the chosen value. Previous work suggests that these cell groups form a decision circuit, but the anatomical organization of this circuit is poorly understood. Using calcium imaging, we recorded from layer 2/3 (L2/3) and layer 5 (L5) of mice choosing between juice flavors. Decision variables were differentially represented across layers: juice-specific offer values and their spatial configuration were predominant in L2/3, while spatial offer values, chosen side, and chosen value were predominant in L5. Within each layer, functional cell groups were organized in clusters. The temporal dynamics of neural signals in the two layers indicated a combination of feed-forward and feed-back processes, and pointed to L5 as the locus for winner-take-all value comparison. These results reveal that economic decisions rely on a complex architecture distributed across layers of OFC.

## Introduction

Economic choice behavior entails representing and comparing the subjective values of two (or more) offers. These operations are known to depend on OFC^1–3^. Stimulation studies in non-human primates showed that offer values represented in this area are causally related to choices^4^, and that neuronal activity in OFC contributes to value comparison^5^. Concurrently, research in humans^6, 7^ and rodents^8–10^ found that OFC lesion or inactivation selectively disrupts choices. Neurophysiology experiments examined the activity of individual cells while animals chose between juice flavors offered in variable quantities. Different groups of neurons were found to encode different variables, including individual offer values, the binary choice outcome, and the chosen value^9–14^. Notably, these variables capture both the input (offer value) and the output (choice outcome, chosen value) of the decision process, suggesting that the cell groups identified in OFC constitute the building blocks of a decision circuit. Supporting this hypothesis, trial-to-trial fluctuations in the activity of each cell group correlates with variability in choices^12, 15–17^. Moreover, modeling work indicated that the cell groups found in OFC are computationally sufficient to generate binary decisions^18–23^. Importantly, the anatomical organization of this neural circuit is poorly understood. A circuit-level understanding of economic decisions would pave the way to treat mental disorders affecting choices.

Here we examined whether variables encoded in OFC are differentially distributed across cortical layers. Using two-photon (2P) calcium (Ca^2+^) imaging, we recorded neuronal activity in the OFC of mice engaged in a juice-choice task. The mouse OFC lacks layer 4. Thus cortical and subcortical afferent connections reach OFC primarily in L2/3, which can be regarded as the input layer of this area. Conversely, efferent projections from OFC originate in both L2/3 and L5^24, 25^. Thus at the outset of this study, we hypothesized that neurons encoding the offer values might preferentially populate L2/3. A weaker hypothesis was that neurons encoding the choice outcome and the chosen value might preferentially populate L5. Our results were broadly consistent with this picture, but also revealed a more complex architecture. Neurons in OFC encoded juice-specific offer values, their spatial configuration, spatial offer values, the binary choice outcome, and the chosen value. The representation of these variables was not strictly segregated to specific cortical layers – i.e., all variables were represented in each layer. However, the strength of these neural signals and their temporal dynamics differed significantly across layers. Our analyses indicated that L2/3 cells encoding juice-specific offer values and their spatial configuration informed L5 cells encoding spatial offer values; that the winner-take-all process resulting in a binary choice outcome took place within L5; and that L5 sent feedback signals representing the choice outcome to L2/3. This emerging picture was consistent with known patterns of anatomical connectivity^24–28^ and with computational considerations. It was also confirmed by estimates of functional connectivity obtained using Granger causality analysis (GCA)^29–31^. Taken together, our results provide strong evidence for a laminar organization of the decision circuit within OFC and for a primary role of L5 in winner-take-all value comparison.

## Results

### Economic choices in mice

During the experiments, animals were head-restrained and placed under a microscope (**Fig.1**). Mice were trained to perform a juice-choice task (**Fig.1B**). In each session, the animal chose between two juice flavors labeled A and B (with A preferred) offered in variable quantities. The task closely resembled that used in monkey studies^11^, except that offers were represented by olfactory stimuli. The odor identity (octanal or octanol) was associated with the juice flavor while the odor concentration indicated the juice quantity. For each juice flavor, we used 4-5 quantity levels. The two odors were presented on the two sides of the animal’s nose, and the mouse indicated its choice by licking one of two spouts placed near its mouth. Offered quantities varied from trial to trial and the spatial configuration of the offers was counterbalanced across trials in each session (see **Methods**).

**Figure 1.**
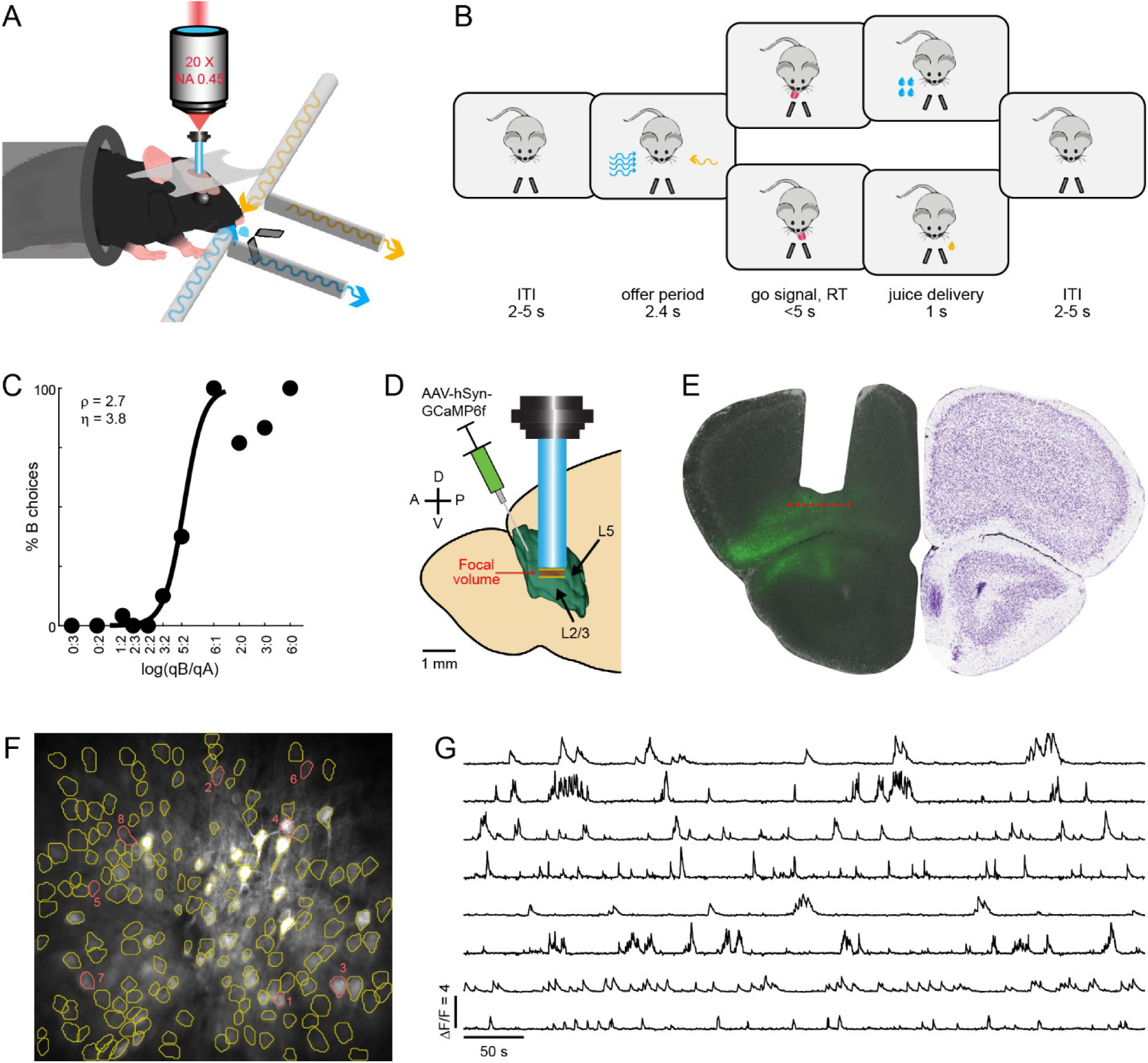
Choice task and neural recordings in OFC. **A.** Apparatus. The animal was head-fixed, restrained inside a tube, and placed under the 2P microscope. Olfactory stimuli were delivered through two odor ports (left and right side), and two additional ports vacuumed the air before each trial. Two reward spouts (left and right side) detected the animal’s choice (first lick following the go signal) and delivered the chosen juice. **B.** Choice task. Following the intertrial interval (ITI), the animal was presented two odors in variable concentration. Each odor was associated with a juice flavor and the concentration represented the offered quantity. After the offer period (2.4-2.8 s), a tone indicated the go signal. The animal had to respond with a lick to the left or right spout within 5 s, and the chosen juice was delivered immediately thereafter. The spatial configuration of the offers (A/B left/right, or vice versa) varied from trial to trial pseudo randomly. **C.** Example session. The x-axis represents the log quantity ratio log(*q_B_*/*q_A_*), where *q_A_* and *q_B_* were the offered quantities. The y-axis represents the percent of trials in which the animal chose juice B. The black line was derived from a logistic regression, which provided measures for the relative value (*ρ*) and the choice accuracy (*η*). **D.** Surgical preparation. To perform *in vivo* Ca^2+^ imaging, we prepared animals to express GCaMP6f in OFC and we implanted a GRIN lens. The cartoon illustrates the anterior part of the mouse brain from a sagittal view. The green area represents the mouse orbitofrontal area and its layers (L2/3, L5). The mouse OFC lacks layer 4. A, anterior, P, posterior, D, dorsal, V, ventral. **E.** Histology. The location of each GRIN lens was reconstructed at the end of the experiments. Shown here are an example animal from our experiments (left) and an image from the Allen Brain Atlas (right). On the left, gray is DAPI; green is GCaMP6 expression; the missing cortex shows the location of the GRIN lens. **F.** Example field of view. The gray signal is GCaMP6f, averaged across 8 frames for visualization purposes. The cell segmentation was performed using the entire session. Each yellow or red circle represents one neuron. **G.** ΔF/F signals. Each trace shows the ΔF/F in the first 5000 frames of the session, for each of the eight cells in red in panel F (numbered top to bottom). The activity was acquired while the animal performed choices and it is aligned here at the start of the session.

Choices reliably presented a quality-quantity trade-off between juice flavor and juice quantity (**Fig.1C**). In each session, choice data were submitted to a logistic analysis, which provided measures for the relative value of the juices (*ρ*) and the choice accuracy (*ƞ*; see **Methods**). Intuitively, indicating with *q_A_* and *q_B_* the juice quantities offered in each trial, *ρ* was the quantity ratio *q_B_*/*q_A_* that made the animal indifferent between the two offers; *ƞ* was proportional to the sigmoid steepness and inversely related to choice variability^32^. As in previous studies, *ρ* measured in a particular session was used to compute the variables possibly encoded by individual neurons recorded in the same session^11, 14^.

### Representation of decision variables in OFC

Experiments were conducted in 14 mice prepared through viral injection to express GCaMP6f specifically in OFC (**Suppl. Table 1**, **Suppl. Table 2**). 2P Ca^2+^ imaging was performed through a Gradient-Index (GRIN) lens (1 mm diameter, 4 mm length) implanted above OFC. Fields of view (FOVs) were located 50-350 µm below the GRIN lens (≥20 µm distance between FOVs; 3-12 FOVs per animal). Ca^2+^ activity was processed in Matlab using the CaImAn package^33^. For each neuron, fluorescence traces were normalized by calculating ΔF/F (**Fig.1FG**). Neural data included in this study came from 78 FOVs. In any FOV, we could identify 18-225 cells (mean = 93, median = 89). Thus our entire data set included 7261 individual cells (see **Methods**).

Neuronal activity was analyzed in five time windows aligned with offer onset and juice delivery onset (see **Methods**). A “trial type” was defined by two offered quantities, their spatial configuration, and a choice. For each cell, each time window, and each trial type, we averaged ΔF/F signals over trials. A neuronal response was defined as the activity of one neuron in one time window as a function of the trial type. As previously described^14^, individual cells in OFC represented different decision variables, including offer values in a good-based reference frame (**Fig.2A-C**), the spatial configuration of the offers (**Fig.2D**), offer values in a spatial reference frame (**Fig.2EF**), the chosen value (**Fig.2G**) and the chosen side (**Fig.2H**). Each of these variables could be encoded with positive or negative slope.

**Figure 2.**
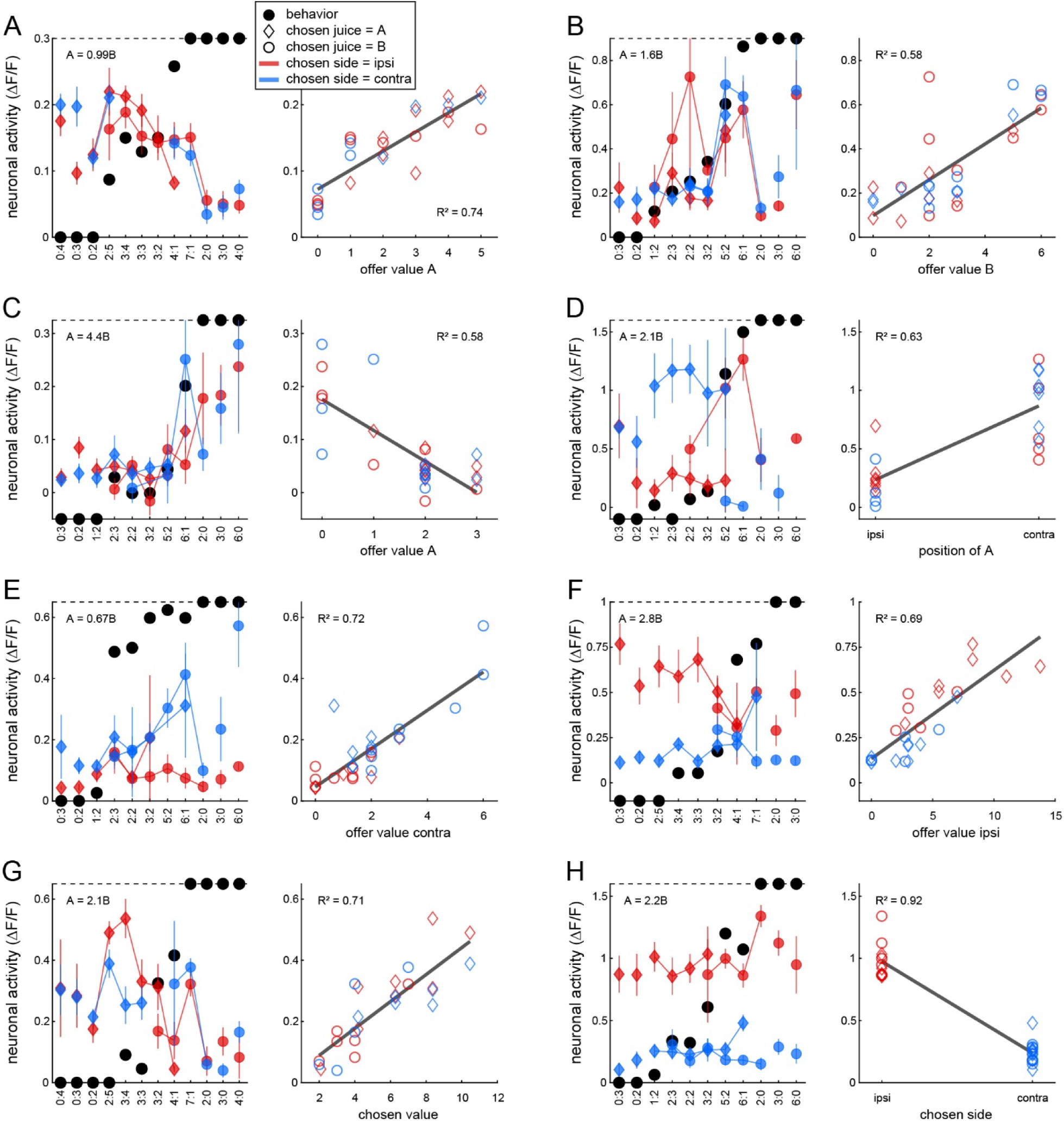
OFC, example cells. **A.** Neuronal response encoding the *offer value A* (L2/3, late delay). In the left panel, the x-axis represents different offer types ranked by the quantity ratio q_B_/q_A_. Black dots represent the choice pattern – i.e., for each offer type, the fraction of trials in which the animal chose juice B. Color symbols represent neural activity (ΔF/F), with each data point representing one trial type. Trial in which the animal chose left (ipsi) and right (contra) are plotted in red and blue, respectively. Trials in which the animal chose A and B are represented by diamonds and circles, respectively. Error bars indicate s.e. In the right panel, the same neuronal response is plotted against variable *offer value A*. The gray line is from a linear regression, and the R^2^ is indicated. **B.** Neuronal response encoding the *offer value B* (L2/3, post-offer). **C.** Neuronal response encoding the *offer value A* with negative slope (L5, pre-choice). **D.** Neuronal response encoding the *position of A* (L2/3 pre-choice). **E.** Neuronal response encoding the *offer value C* (L5, pre-choice). **F.** Neuronal response encoding the *offer value I* (L5, pre-choice). **G.** Neuronal response encoding the *chosen value* (L2/3, pre-choice). **H.** Neuronal response encoding the *chosen side* (L5, post-choice).

We conducted a series of analyses to confirm these observations. First, the activity of each cell in each time window was submitted to a 1-way ANOVA (factor: trial type). Neurons that passed the significance threshold (p<0.01) in ≥1 time window were identified as task-related and included in subsequent analyses (N = 1804 cells; **Suppl. Table 3**). Second, we defined a series of candidate variables that neurons in OFC might potentially encode (**Suppl. Table 4**). For each response passing the ANOVA criterion, we performed a linear regression against each variable. If the regression slope differed significantly from zero (p<0.05), the variable was said to explain the response. For each variable, the regression also provided the R^2^. For variables that did not explain the neuronal response, we set R^2^ = 0. Third, for each response passing the ANOVA criterion, we identified the variable that provided the best fit (highest R^2^), and we thus constructed a population table. For each time window, we then computed the number of responses best explained by each variable (**Suppl. Fig.1A-C**). Fourth, we used a best-subset procedure to identify a smaller set of variables that would account for the whole neuronal population. This procedure identified variables *position of A*, *offer value A*, *offer value B*, *offer value I* (*I* = *ipsilateral*), *offer value C* (*C* = *contralateral*), *chosen side*, and *chosen value*, which collectively explained 81% of task-related responses (**Suppl. Fig.1DE**). Fifth, we conducted a post-hoc analysis to compare the explanatory power of selected variables with that of other candidate variables. We found that the explanatory power of all selected variables was significantly higher (**Suppl. Table 5**). On these bases, we classified each cell by assigning it to the variable providing the best explanation across time windows (largest total R^2^; **Suppl. Fig.2**; see **Methods**). Cells that did not pass the ANOVA criterion in any time window and task-related cells that were not explained by any variable were collectively classified as *untuned*.

### Visual classification of neurons in L2/3 and L5

As illustrated in **Fig.1D**, cortical layers in OFC were not parallel to the focal plane. Thus, depending on the depth of the GRIN lens and the focal distance of the FOV, the imaging plane could lie within L2/3, include both L2/3 and L5, or lie within L5. Across FOVs, we identified substantial and reliable differences between cells from different layers. Cells in L2/3 were smaller, brighter, more round, often spiny, and more densely packed (estimated density: 40-50 cells/(100 μm)^3^). Conversely, cells in L5 were larger, dimmer, and more sparsely packed (estimated density: 8-12 cells/(100 μm)^3^). Most revealingly, L5 cells showed pyramidal cell bodies with spine-covered apical dendrites oriented anteriorly towards superficial layers. One example FOV that crossed both layers illustrates these characteristics (**Fig.3AB; Suppl. Fig.3**). In all our animals, small cells (L2/3) were located more anterior and more ventral, while large cells (L5) were located more posterior and more dorsal, consistent with the relative orientations of cortical layers and the GRIN lens.

**Figure 3.**
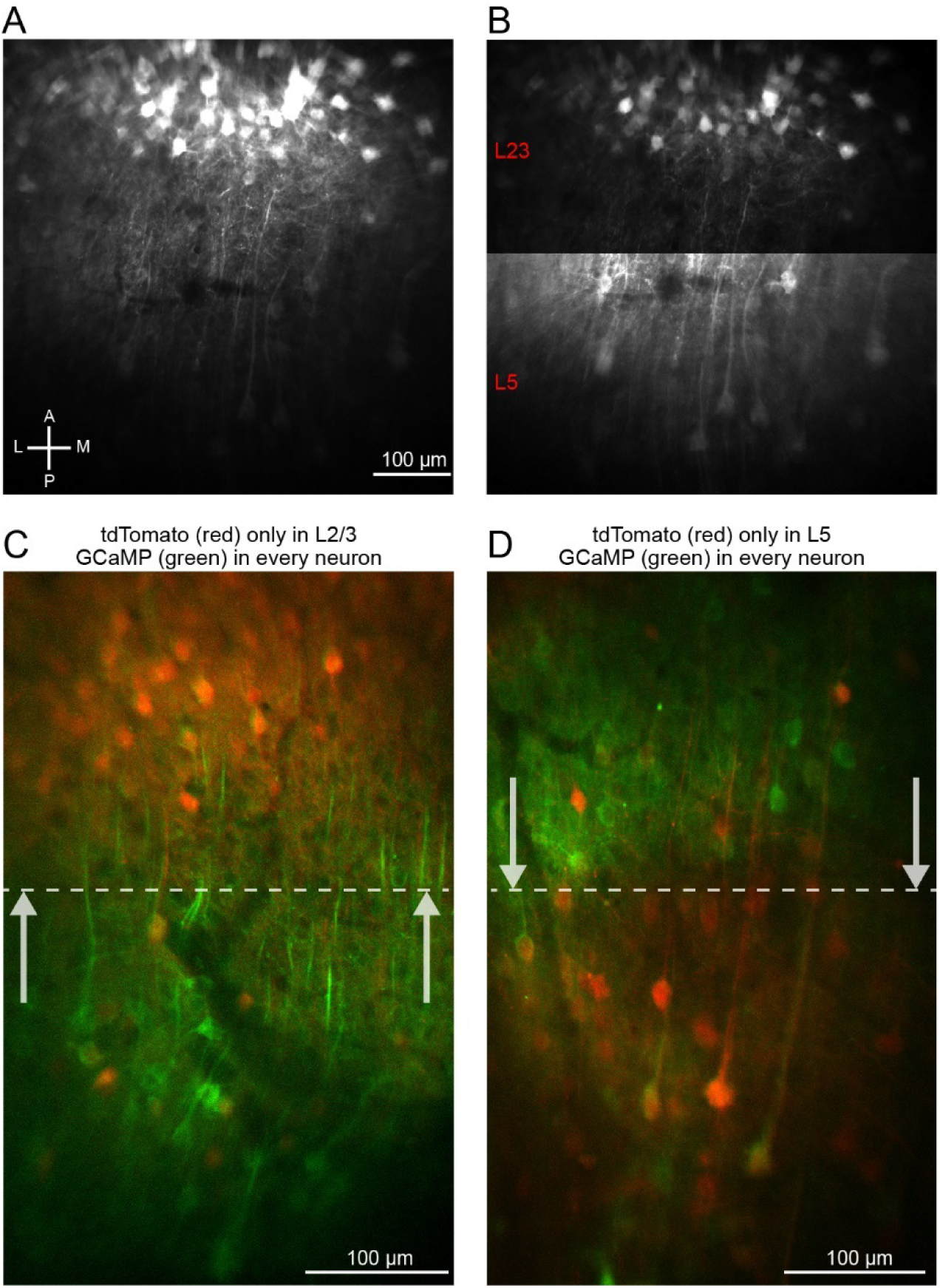
In vivo imaging of different cortical layers. **A.** Example FOV with two distinct bands of neurons. Using GCaMP signal, it is possible to discriminate two laminar bands of cells with different anatomical and morphological features. The first band lays more superficial and anterior (upper part of this section); the second band lays more deep and posterior (lower part of this section). A, anterior, P, posterior, L, lateral, M, medial. **B.** Improved visualization. Same FOV as in panel A, but the contrast was independently adjusted for the upper and lower parts of the section, to better visualize the two cell bands. In the upper part, neurons were smaller, brighter, and more dense; in the lower part, neurons were larger, dimmer, less dense, and presented an apical dendrite directed towards layer 1. **C.** Identification of small cells as L2/3 cells. This experiment was conducted in a transgenic mouse expressing Cre only in L2/3 (Rasgrf2-2A-dCre). Viral expression was achieved by injecting a cocktail of two viruses to express GCaMP6 constitutively (i.e., in every neuron; green signal) and tdTomato in Cre-dependent way (i.e., only in L2/3; red signal) (see **Methods**). Notably, tdTomato only labeled cells with morphological features similar to those observed in the upper part of panels AB. **D.** Identification of large cells as L5 cells. This experiment was conducted in a transgenic mouse expressing Cre only in L5 (ER81-CreERT2). In this case, tdTomato labeled cells with morphological features similar to those observed in the lower part of panels AB. Dashed lines in panels CD indicate the putative border between L2/3 and L5.

We confirmed the separability between small and large cells with two distinct approaches. First, on a subset of FOVs we verified that we could accurately classify cells with an unsupervised algorithm based purely on cellular morphology (**Suppl. Fig.4; Methods**). Second, to unambiguously validate the identification of small cells with L2/3 and large cells with L5, we used two strains of mice that expressed Cre only in L2/3 (Rasgrf2-2A-dCre) or only in L5 (ER81-CreERT2). In both strains, we used viral injection to express GCaMP6f in all neurons and tdTomato in a Cre-dependent way (**Suppl. Fig.5; Suppl. Table 2**; **Methods**). As illustrated in **Fig.3CD**, these experiments confirmed our interpretation. In animals expressing tdTomato in L2/3, only small cells were labeled in red; conversely, in animals expressing tdTomato in L5, only large cells were labeled in red.

**Figure 4.**
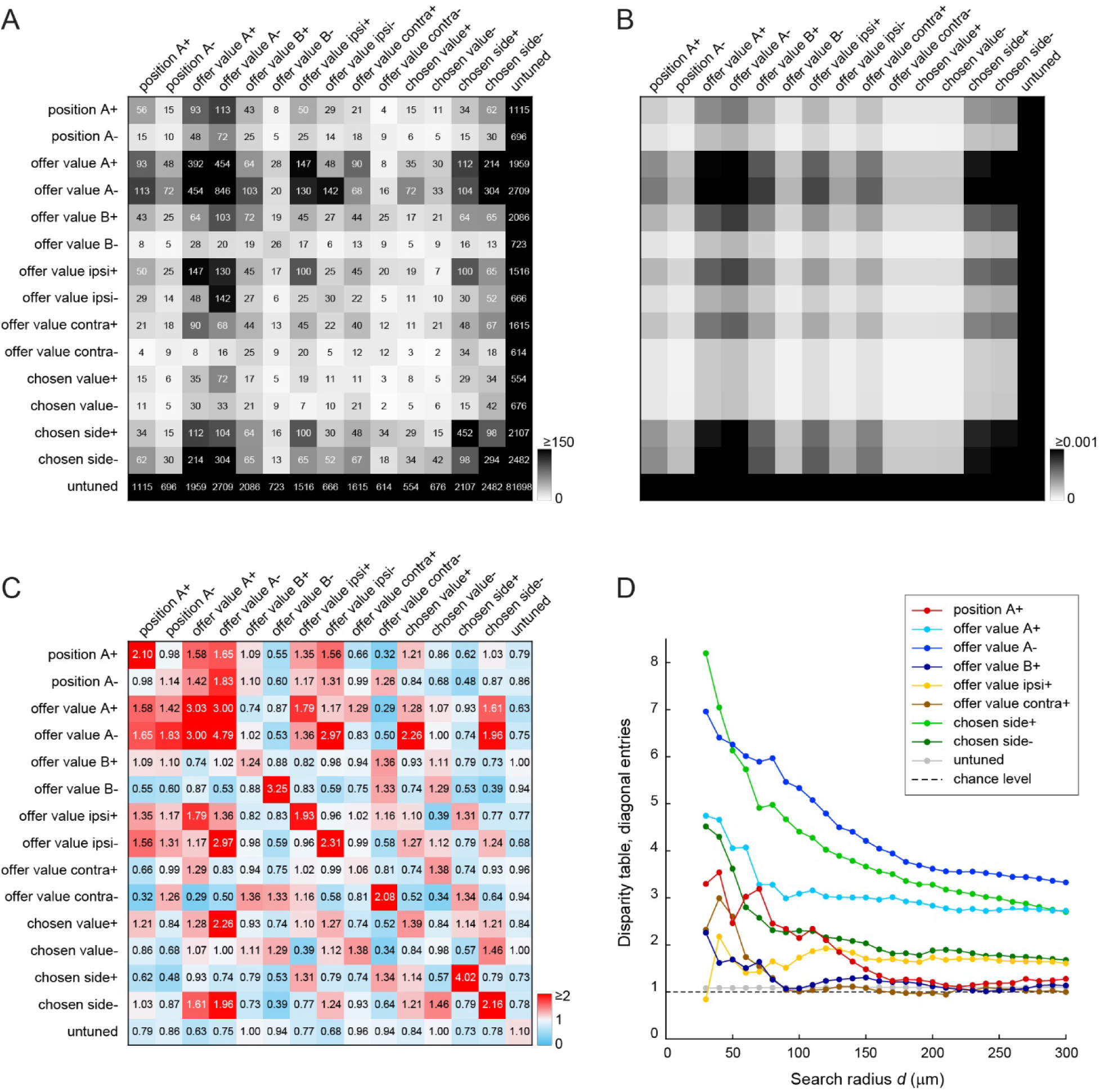
Spatial clustering of functional cell groups in L2/3. **A.** Table of cell counts (search radius *d* = 120 μm). Rows and columns corresponding to variables encoded by the reference cell and by cells in the vicinity of the reference cell, respectively. Shades of gray recapitulate cell counts (see color bar). Of note, the table of cell counts was symmetric because each pair of cells (*i*, *j*) contributed twice – once with *i* as the reference cell (row) and *j* as a cell in the vicinity of *i* (column), and once with *i* and *j* in reversed roles. **B.** Table of expected frequencies. In this table, entry (*u*, *v*) is the probability that two cells randomly selected encode variables *u* and *v*. This table is derived from the frequencies shown in **Suppl.** Fig.2. **C.** Disparity table. This table was computed by normalizing cell counts (panel A) and dividing them by the expected frequencies (panel B). For all entries, chance level is 1 (white); entries <1 (blue) or >1 (red) indicate that cell counts were smaller or larger than expected by chance, respectively. **D.** Diagonal entries of the disparity table as a function of the search radius (*d*). Each line and color represents a particular variable (see legend). Only variables *u* with expected frequency p*_uu_* ≥ 0.0002 were included (see **Methods**).

**Figure 5.**
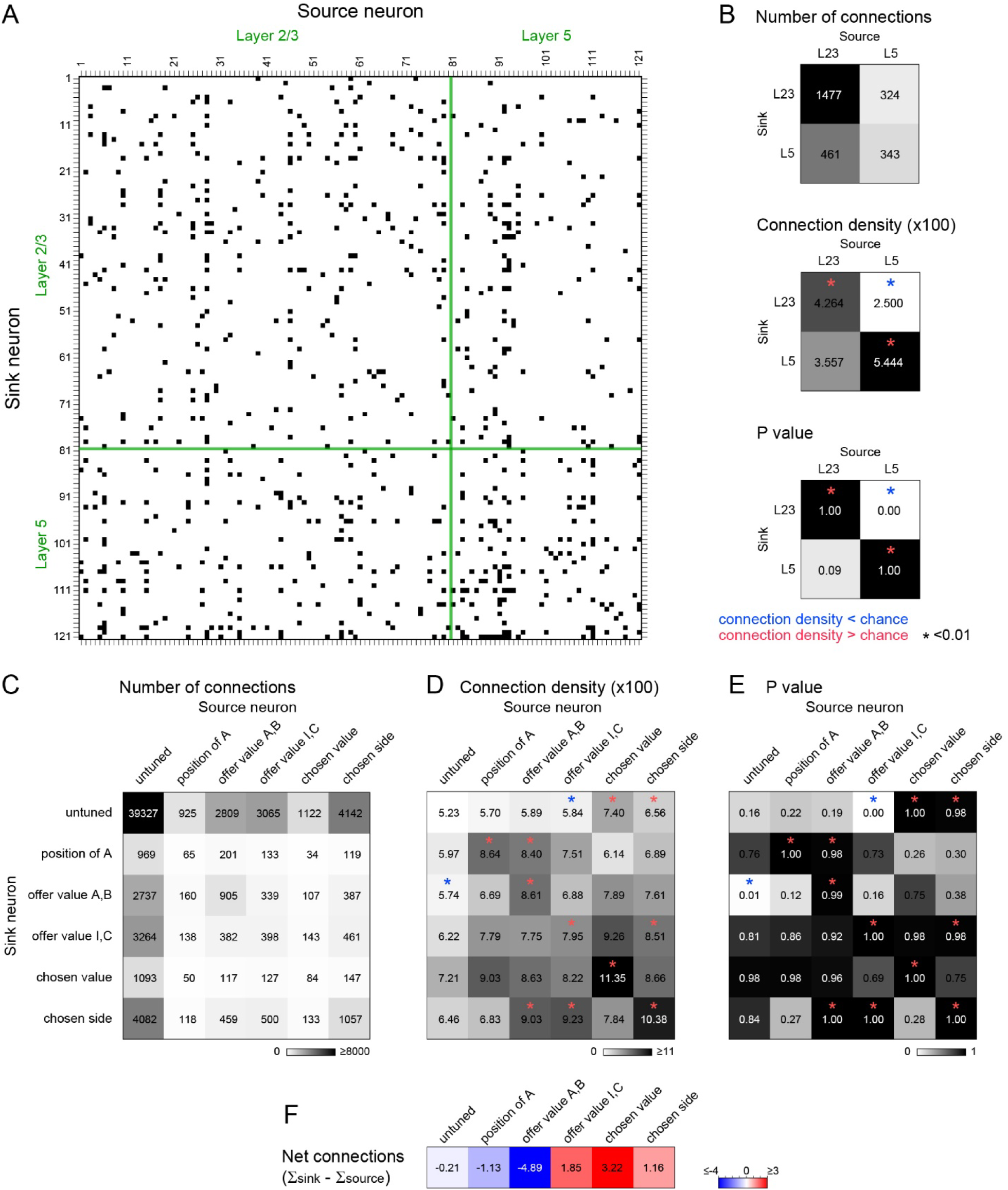
Functional connectivity of the decision circuit. The figure illustrates the results of Granger causality analysis (GCA). **A.** Example FOV. The FOV crossed layers and included N = 121 cells (80 from L2/3; 41 from L5). In the connectivity matrix shown here, columns and rows represent source neurons and sink neurons, respectively. Black squares indicate significant GCA connections (p< 0.01). Cells were sorted such to divide the connectivity matrix in four sectors corresponding to L2/3→L2/3, L2/3→L5, L5→L2/3, and L5→L5 connections. **B.** Connectivity within and between cortical layers. We pooled 11 FOVs that crossed L2/3 and L5 (see main text). Entries in the center panel are mean connection densities (x100, for readability) for the 4 sectors. Asterisks indicate significant departures from chance level, detected using a bootstrap analysis (blue for density below chance; red for density above chance). **CDE.** Connectivity within and between functional groups. Neurons were classified in 6 functional groups (*position of A*, *offer value AB*, *offer value IC*, *chosen side*, *chosen value*, *untuned*). Thus for each FOV, the connectivity matrix was divided in 36 sectors. For a population analysis, we pooled data from both layers. Entries in the matrix shown here are mean connection densities (x100, for readability) for the 36 sectors. Asterisks indicate significant departures from chance level (bootstrap analysis). **F.** Net connectivity. For each cell group, we computed the difference between the total connection density as sink and the total connection density as a source. Hence, values <0 and >0 indicate the cell group was primarily a source or a sink of connections, respectively.

On these bases, we classified every cell in our data set visually as L2/3 or L5. This classification was conducted independently of any other analysis. The resulting data set included 3134 cells from L2/3 and 2027 cells from L5. An additional 2100 cells could not be reliably classified (see **Methods**).

### Functional cell groups form anatomical clusters

We next investigated whether neurons encoding any particular variable were spatially clustered. We examined the two layers separately, starting with L2/3. We set a minimum distance *d_0_* = 20 μm and a search radius *d* = 120 μm. For each FOV and for each neuron *i*, we examined cells in the vicinity of *i* – i.e., cells whose distance from *i* was larger than *d_0_* but smaller than *d* (see **Methods, Suppl. Fig.6**). We constructed a table where rows indicated the variable encoded by neuron *i*, columns indicated the variables encoded by cells in the vicinity of *i*, and entries were cell counts. We repeated this operation for each neuron and each FOV, and we summed the resulting tables. Thus we obtained a table of cells counts for the whole data set (**Fig.4A**).

**Figure 6.**
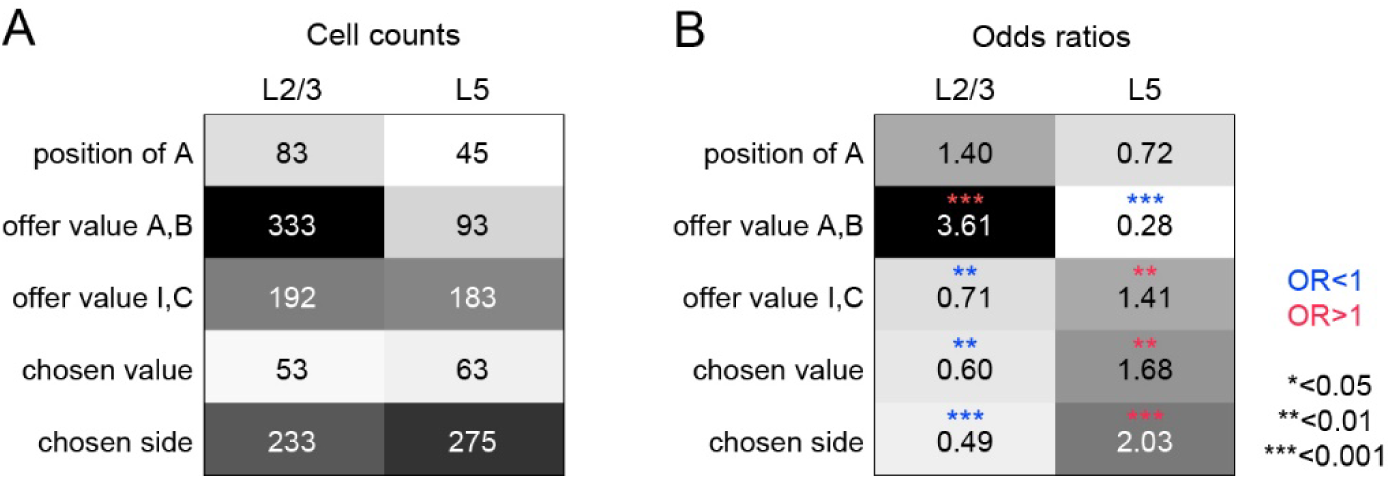
Different decision variables are preferentially represented in L2/3 and L5. **A.** Table of cell counts. For each cortical layer (column), entries in this table indicate the number of cells classified as encoding each variable (rows). The gray scale recapitulates cell counts. **B.** Odds ratios. Each entry indicates the OR computed from the contingency table (panel A). Asterisks (*) indicate significant departure from chance (blue for OR<1; red for OR>1). Exact p values: p=0.09 (*position of A*), p<10^-24^ (*offer value AB*), p<0.005 (*offer value IC*), p<10^-10^ (*chosen value*), p<0.009 (*chosen side*; Fisher’s exact test).

Entries in the table of cell counts differed substantially depending on the pair of variables. This fact was not surprising because each variable was encoded by different fractions of cells (**Suppl. Fig.2A**). We aimed to assess whether entries in this table – especially those on the diagonal – were higher than expected by chance. To do so, we normalized the table of cell counts. We also computed a table of expected frequencies based on the fraction of cells encoding each variable across the entire population (**Fig.4B**). Finally, we defined the disparity table as the entry-by-entry ratio between the table of normalized cell counts and the table of expected frequencies (**Fig.4C**). For each entry in the disparity table, chance level = 1; entries <1 or >1 indicated that the corresponding cell count was below or above chance, respectively.

Three aspects of **Fig.4C** are notable. First, all non-diagonal entries in the rightmost column were ≤1. This observation suggests some degree of segregation between tuned and untuned cells. Second, 14 of 15 entries on the main diagonal were >1. In other words, the likelihood of finding two cells encoding the same variable in the vicinity of each other exceeded chance level for all but one variable. This observation suggests some degree of clustering for cells encoding any one variable. Third, non-diagonal entries in the table were not all <1. In other words, spatial clustering was not rigid. We repeated this analysis varying the search radius *d* between 30 μm and 300 μm, and we assessed how diagonal entries in the disparity table varied as a function of *d*. If neurons encoding a particular variable were spatially clustered, the corresponding diagonal entry in the disparity table should be always >1 and decrease with *d*. This is indeed what we found for each variable (**Fig.4D**; see **Methods**).

We repeated this analysis for L5 and we obtained a qualitatively similar picture (**Suppl. Fig.7**). Control analyses where we doubled *d_0_* confirmed these results (**Suppl. Fig.8**). In summary, neurons encoding any specific variable presented some degree of anatomical clustering in both L2/3 and L5. Inspection of **Fig.4D** and **Suppl. Fig.7D** suggests a cluster diameter on the order of 100 μm. Interestingly, we observed clusters of the same scale in the myelination patterns of superficial layer 1 (L1; **Suppl. Fig.9**).

**Figure 7.**
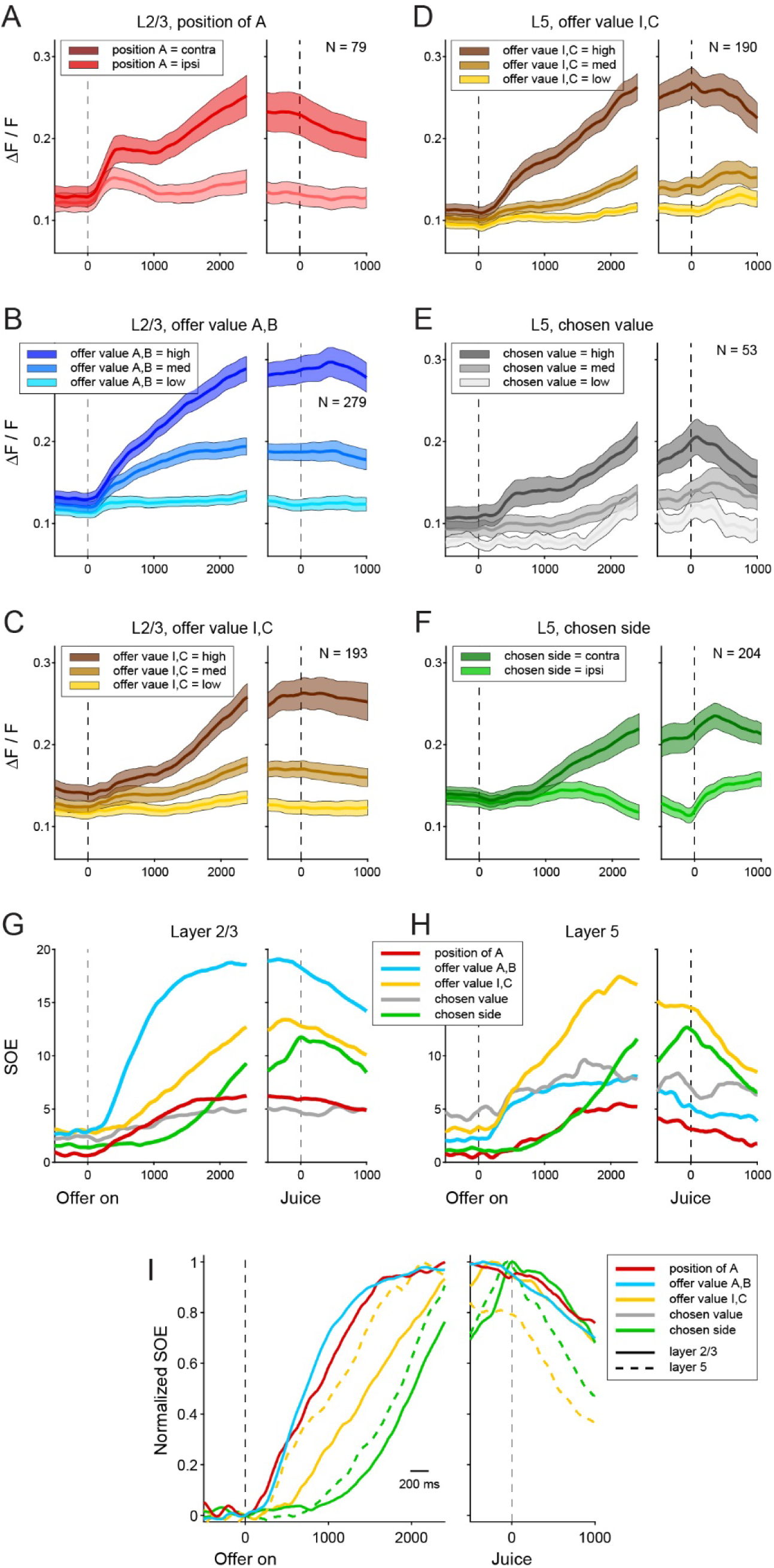
Activity profiles and strength of the encoding. **A.** L2/3, cells encoding the *position of A*, activity profiles. For each cell, trials were divided depending on whether juice A was offered on the ipsi or contra side (*position of A* = ipsi or contra). Trials were aligned separately at offer onset and juice delivery. We examined neural signals (ΔF/F), obtained an activity profile for each neuron, and averaged the activity profiles across neurons (see **Methods**). **B.** L2/3, cells encoding the *offer value A,B*. We pooled neurons encoding variables *offer value A* and *offer value B*. For each cell, we divided trials in three groups depending on whether the relevant offer value was low, medium, or high (see **Methods**). We obtained an activity profile for each neuron and for each value bin, and we averaged the activity profiles across neurons for each value bin. **C.** L2/3, cells encoding the *offer value I,C*. **D.** L5, cells encoding the *offer value I,C*. **E.** L5, cells encoding the *chosen value*. **F.** L5, cells encoding the *chosen side*. Panels A-F illustrate only cells with positive encoding (complete data set shown in **Suppl.** Fig.12). In each panel, lines are means, running error bars equal 2 s.e., and the number of cells contributing to the population is indicated. All times are in ms. **GH.** Strength of the encoding (SOE). For each variable, we computed the combined SOE for positive- and negative-encoding cells (see **Methods**). Traces shown here illustrate the combined SOE for each cell group (see color legend), separately for L2/3 (panel G) and L5 (panel H). **I.** Normalized SOE. Traces in panels GH vary both in amplitude and timing. To specifically assess differences in timing, we normalized each trace by subtracting the value at offer onset and by dividing by baseline-subtracted trace by its maximum. Panel I shows the normalized traces from the two layers for the most prominent variables (continuous lines = L2/3, dashed lines = L5). First to emerge after offer onset were variables *offer value AB* and *position of A* in L2/3. These were followed by variable *offer value IC* in L5 and, lastly, by variable *chosen side*. Note that both variables *offer value IC and chosen side* emerged in L5 first and were substantially delayed in L2/3 (by 200-300 ms), suggesting a feedback signal.

**Figure 8.**
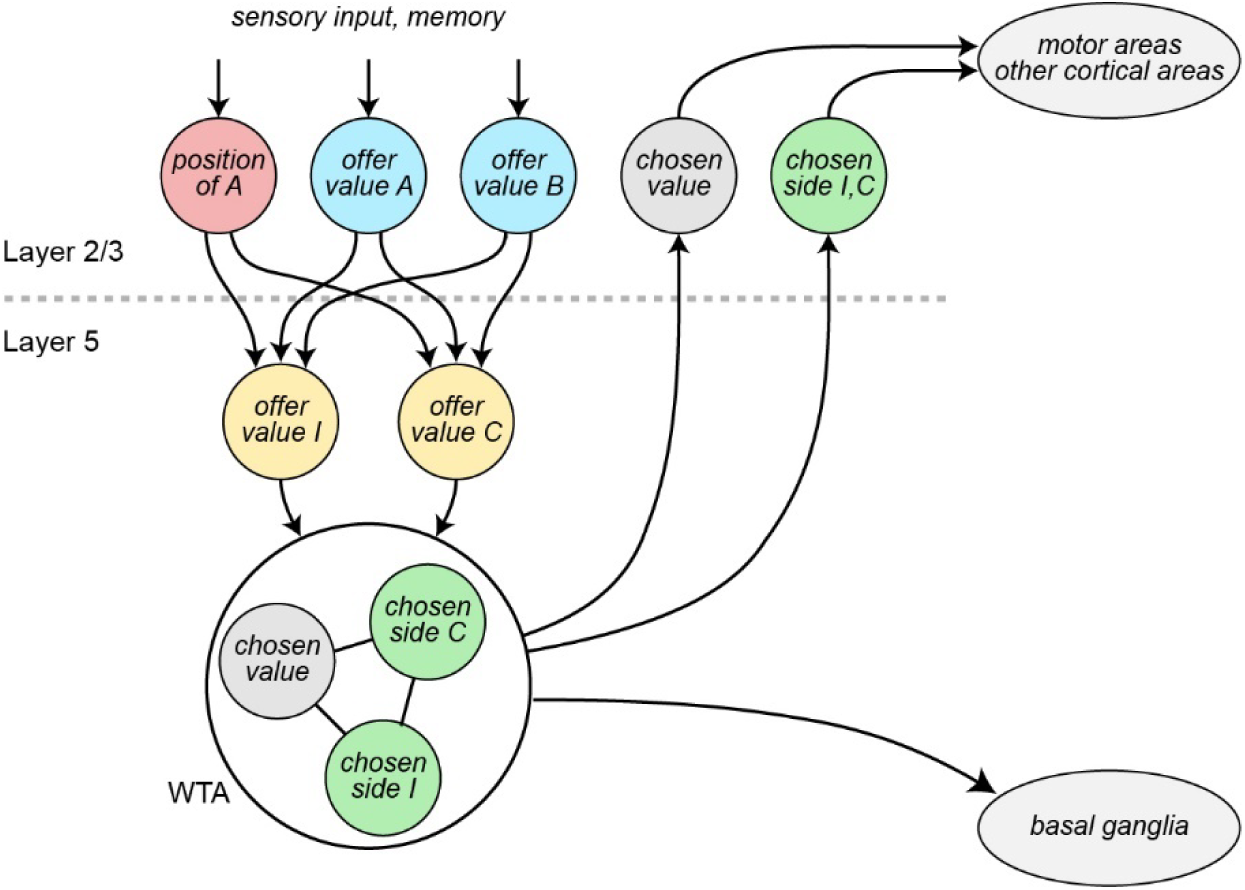
Emerging circuit organization. WTA = winner-take-all.

### Functional connectivity of the decision circuit

In subsequent analyses, *offer value A* and *offer value B* cells were pooled and collectively labeled *offer value AB* cells. Similarly, *offer value I* and *offer value C* cells were collectively labeled *offer value IC* cells.

The variables encoded in OFC capture both the input and the output of the decision process. Furthermore, this neuronal population represents both good-based variables (*offer value AB*) and spatial variables (*offer value IC*, *chosen side*), as well as the *position of A*, which bridges the two reference frames. This observation suggests that the mouse OFC hosts both a good-to-action transformation and a winner-take-all process of value comparison. More specifically, the fact that *offer value IC* is a pre-decision variable and the scarcity of *chosen juice* signals in the mouse OFC suggest that the good-to-action transformation takes place upstream of value comparison. In this view, signals *offer value AB* and *position of A* would be combined to compute *offer value I* and *offer value C*. In turn, these signals would feed the winner-take-all process resulting in *chosen side* and *chosen value*.

We tested these hypotheses using GCA, which provided estimates of functional connectivity between cortical layers and cell groups. In essence, GCA is a measure of temporally directed correlation between time series^29–31^ such as the ΔF/F signals of two neurons^34–36^. Consider two cells X and Y. If the past activity of X provides information about the future activity of Y beyond the information already provided by the past activity of Y, one can say that there is a flow of information from X to Y. In this case, X is said to be functionally connected to Y (X→Y). In the relation X→Y, cell X and cell Y are termed the source and the sink of the connection, respectively. Importantly, X→Y does not imply (or exclude) Y→X.

First, we examined the connectivity within and between cortical layers. This analysis focused on FOVs including both L2/3 and L5 and presenting a clear border between the two layers (11 FOVs from 3 animals). On average, each of these sessions had 76 neurons (median = 69, std = 22). For each FOV, we sorted cells in the connectivity matrix so to obtain 4 sectors corresponding to the 4 connection patterns L2/3→L2/3, L2/3→L5, L5→L2/3, and L5→L5. **Fig.5A** illustrates one example session with N = 121 cells (80 cells in L2/3, 41 cells in L5). For each sector, we obtained the number of statistically significant connections (p<0.01) and the number of possible connections (see **Methods**). For a population analysis, we combined these numbers across the 11 FOVs and we computed the connection density for each of the 4 sectors (**Fig.5B**). Finally, for each sector we used a bootstrap analysis to assess whether the connection density differed significantly from that expected by chance (p<0.05). Two aspects of the results were most noteworthy. First, the density of connections within layers was generally higher than that between layers. Second, focusing on the connectivity between layers, the density of L2/3→L5 connections was higher than the density of L5→L2/3 connections. Importantly, these two observations recapitulate known anatomical facts – i.e., that the anatomical connectivity within layers is higher than that between layers, and that anatomical projections from L2/3 to L5 are stronger than projections from L5 to L2/3^24, 25^. Thus the results of **Fig.5B** can be viewed as validating the GCA approach.

Next, we examined the connectivity between groups of neurons encoding different variables. For this analysis, we pooled FOVs from different layers. We considered the 6 cell groups *position of A*, *offer value AB*, *offer value IC*, *chosen side*, *chosen value*, and *untuned*. Thus data from each session could be divided in 36 sectors. For each FOV and each sector, we computed the number of statistically significant connections (p<0.01) and the number of possible connections. We then pooled the results from all FOVs and we computed the connection density for each of the 36 sectors (**Fig.5CD**). For each entry in the table, a bootstrap analysis assessed whether the connection density differed significantly from chance (**Fig.5E**). Two results were most noteworthy.

First, for all entries on the diagonal, the density of connections was significantly higher than chance. In other words, cells encoding any variable were preferentially connected to other cells encoding the same variable. This observation is consistent with anatomical studies showing that synaptic connections are denser and stronger when two cells are closer and/or have similar tuning^27, 28, 37, 38^.

Second, there was an overall flow of functional connections directed from *position of A* and *offer value AB* cells to *chosen value IC*, *chosen value*, and *chosen side* cells. This point can be best appreciated by computing, for each cell group, the net connectivity defined as the difference between the total number of incoming connections (sink) and the total number of outgoing connections (source). The net connectivity was <0 for *position of A* and *offer value AB* cells; it was >0 for *chosen value IC*, *chosen value*, and *chosen side* cells (**Fig.5F**). This result supports the notion of a decision circuit in which neurons encoding juice-specific offer values (*offer value AB*) and their spatial configuration (*position of A*) are the primary input, neurons encoding spatial offer values (*offer value IC*) are an intermediate stage, and neurons encoding the choice outcome (*chosen side*, *chosen value*) are the primary output.

### Laminar organization and temporal dynamics of the decision circuit

Finally, we examined how different groups of neurons encoding different variables were distributed across cortical layers. We constructed a contingency table where rows represented encoded variables, columns represented cortical layers, and entries were cell counts (**Fig.6A**). Since different numbers of cells were recorded in L2/3 and L5, cell counts could not be compared directly. Thus for each entry in **Fig.6A**, we computed the corresponding odds ratio (OR), which quantified the strength of the association between the encoded variable and the cortical layer (**Fig.6B; Methods**). For each entry in this table, OR = 1 was the chance level; OR < 1 or OR > 1 indicated that the cell count was smaller or larger than expected by chance, respectively. Fisher’s exact test assessed whether departures from chance were statistically significant.

Inspection of **Fig.6** reveals two important points. First, neurons encoding different decision variables were not strictly segregated – i.e., each decision variable was represented in each cortical layer. At the same time, there were substantial differences between layers. Specifically, neurons encoding variables *position of A* and *offer value AB* were more frequent in L2/3 than in L5 (although this difference reached significance only for the latter). Conversely, neurons encoding variables *offer value IC*, *chosen side*, and *chosen value* were significantly more frequent in L5 than in L2/3.

Beyond cell counts, we aimed to assess how different neural signals differed in strength and timing across cortical layers. Thus for each cortical layer and for each cell group, we constructed a population activity profile (see **Methods**). Cells with positive and negative encoding were examined separately. **Fig.7** illustrates the activity profiles for 6 cell groups, and the whole data set is shown in **Suppl. Fig.10** and **Suppl. Fig.11**. Inspection of **Fig.7** reveals notable differences between layers and variables. In terms of strength, the most intense signals appear to be *offer value AB* in L2/3 (**Fig.7B**) and *offer value IC* in L5 (**Fig.7D**); in contrast, the *chosen value* signal (**Fig.7E**) seems relatively weak. In terms of timing, the signal *position of A* in L2/3 (**Fig.7A**) seems to emerge first, shortly after the offer onset. Next to emerge are, in order, signals *offer value AB* in L2/3 (**Fig.7B**), *offer value IC* in L5 (**Fig.7D**), and *offer value IC* in L2/3 (**Fig.7C**). Last to emerge is the signal *chosen side* (**Fig.7F**).

To quantify these observations, we computed for each variable the strength of encoding (SOE), which essentially captured the signal-to-noise ratio of each decision variable over the course of the trial (see **Methods**). SOE was computed separately for positive- and negative-encoding cells (**Suppl. Fig.12**) and then combined across the two cell groups for each variable. SOE traces confirmed that neural signals from different layers differed significantly in strength and timing (**Fig.7GH**). To specifically assess differences in timing, we normalized each SOE trace by subtracting the value at offer onset and by dividing the baseline-subtracted trace by its maximum. Inspection of **Fig.7I**, which illustrates the normalized SOE traces from both layers, is most revealing. First to emerge after offer onset were signals *offer value AB* and *position of A* in L2/3. These were followed by variable *offer value IC* and, lastly, by variable *chosen side*. Critically, both variables *offer value IC and chosen side* emerged in L5 substantially earlier than in L2/3, where they materialized with a delay of 200-300 ms. These temporal profiles suggest an initial flow of information from L2/3 to L5, followed by a computation and a winner-take-all process in L5, and by a feedback signal from L5 to L2/3.

## Discussion

The goal of this study was to shed light on the organization of a neural circuit known to play a central role in the formation of economic decisions. Our findings can be interpreted in the light of (1) anatomical patterns of connectivity between brain regions and cortical layers^24, 25^ and (2) computational considerations on how functional cell groups might contribute to the decision process. Anatomically, afferent projections from sensory and limbic regions reach OFC in L2/3; L2/3 projects densely to L5, which sends feedback projections back to L2/3; finally, both L5 and L2/3 send efferent projections – L2/3 to other cortical areas and L5 (mostly) to subcortical regions. Computationally, the variables identified in OFC suggest that good-based offer values (*offer value AB*) and their spatial configuration (*position of A*) are combined to compute spatial offer values (*offer value IC*); these are then compared in a winner-take-all process leading to a choice outcome (*chosen side*, *chosen value*); once computed, choice outcome signals are broadcast to other brain regions, where they guide suitable actions, inform learning, etc. In this framework, the relative signal strengths and the temporal dynamics recorded in L2/3 and L5, together with the functional connectivity between cell groups, reveal a complex architecture distributed across layers. The picture emerging from our analyses is illustrated in **Fig.8**. In essence, our results suggest an initial flow of information from L2/3 to L5; they point to L5 as the locus for winner-take-all value comparison; and they indicate that the choice outcome is sent from L5 to L2/3 as feedback. Following these computations, efferent projections from L2/3 and L5 presumably broadcast the choice outcome and the chosen value to other cortical and subcortical regions. This scheme may be viewed as a useful framework to guide further investigation. In particular, future research should focus on the mechanisms governing the winner-take-all process in L5, including the role of different groups of interneurons.

Our finding that different computations contributing to a decision process take place in L2/3 and L5 of OFC resonates with previous studies showing layer-specific functional roles in other areas. In primary visual cortex, L2/3 neurons compute differences between top-down motor-related and bottom-up visual inputs^39^, while layer 6 neurons control the gain modulation of neurons in upper layers^40^. In the somatosensory cortex, L2/3 horizontal projections can suppress neighboring L2/3 neurons and simultaneously activate L5 neurons. This coordinated modulation generates competition between neighboring domains^41^. In barrel cortex, L2/3 activity predominantly suppresses L5 activity, enhancing stimulus selectivity and expanding the range of receptive fields^42^. In primary motor cortex, projections from L2/3 to L5 are fractionated depending on whether target neurons are corticospinal or corticostriatal^43^. Furthermore, over the course of motor learning, neurons in L2/3 maintain a stable representation, while neurons in L5a become increasingly aligned with the motor output^44^. Finally, in human lateral prefrontal cortex, superficial layers are preferentially active during working memory tasks, while deeper layers are more active during response periods^45^. Whether these results can be accounted for by unitary principles remains an open question.

An exciting aspect of our results is that neurons in L2/3 and L5 of OFC were found to form functionally homogeneous clusters of ∼100 μm – a finding matched by myelination patterns in L1. Much of cortex is organized in minicolumns or presents some form of functional clustering. However, evidence in this sense is weaker for prefrontal areas, including OFC. In fact, a previous neurophysiology study in monkeys had not revealed any functional clustering of decision variables^46^. Conversely, functional clusters of similar size with persistent activity ahead of upcoming movements have been found in L2/3 of the mouse anterolateral motor cortex^47^. Importantly, our observation builds on the premise that different groups of cells in OFC encode categorically distinct decision variables^12, 48, 49^. Other prefrontal areas showing extensive mixed selectivity might not present similar clustering. Importantly, our clustering analyses were conducted within each cortical layer. Thus the possible presence of columnar organization in OFC remains an open question.

One caveat of this study is that our ability to classify cells in functional groups is not perfect – we previously estimated our success rate at 55% (based on a single session)^14^. This limit likely reflects a combination of factors, including that choices in mice are less accurate than in monkeys, that mice typically complete fewer trials in each session, and that the mouse OFC represents a larger number of variables. To mitigate this issue, we leveraged the fact that this neuronal representation is longitudinally stable^14^. Thus, whenever possible, we classified neurons based on two separate sessions. Critically, remaining classification errors would not easily explain the main results of this study. Indeed, differences in signal strength and temporal dynamics found between cell groups and cortical layers would almost certainly be enhanced, not reduced, if the neuronal classification was entirely error free. Similarly, spatial clustering would almost certainly be enhanced in the absence of any classification error.

With respect to the variables encoded in OFC, our results extend earlier findings. Specifically, a previous study in mice found neurons encoding *position of A*, *offer value IC*, *chosen side*, and *chosen value*^9^ – but not *offer value AB*. This discrepancy might be because the earlier study was based on neurophysiology, with electrodes entering OFC dorsally (i.e., from deep layers). As a consequence, we suspect that neurons in that data set were disproportionally from L5, where *offer value AB* signals are weaker. That aside, it is interesting to reflect on the differences between the two species. In monkeys, the representation of decision variables is good-based, value comparison takes place upstream of the good-to-action transformation, and these two operations appear segregated to different areas (OFC and lateral prefrontal cortex, respectively)^50^. In contrast, in mice, the good-to-action transformation is upstream of value comparison, and both operations take place in OFC. In principle, a design that decouples value comparison from action planning – as observed in monkeys – affords higher flexibility. Thus, good-based decisions might be advantageous, if computationally costly. At the same time, the decision task could be solved entirely in a spatial reference frame. In this perspective, the advantages of adopting two reference frames in the same circuit – as observed here in mice – are unclear. Future research should examine this intriguing question.

## Methods

The present study presents new analyses of previously published data^14^ and new data. All experimental procedures conformed to the NIH *Guide for the Care and Use of Laboratory Animals* and were approved by the Institutional Animal Care and Use Committee (IACUC) at Washington University in St Louis. Unless otherwise noted, all the methods for surgery, viral injection, animal training, behavioral control, neural recordings, image processing, and preliminary data analyses are as detailed in the previous study^14^.

### Animal subjects and surgical procedures

<u>Animal subjects</u>. Data were collected in 19 mice (9 males, 10 females; **Suppl. Table 1**). Of these, 14 mice were used for behavioral and neurophysiology experiments, 2 animals (T01 and T02) were used to generate **Fig.3**, and 3 animals were used only for anatomy experiments (**Suppl. Fig.9**). Animals were housed individually, and behavioral experiments were conducted in the light phase of a 12 hr light/dark cycle. Mice were under water restriction and typically consumed water during the behavioral testing. However, their weight was monitored every testing day to ensure that at least 0.6 mL of liquid reward was consumed^51^ and that the animal maintained their weight ≥75% of the initial record. If either condition was not met, the animal was supplemented with extra water to fulfil the criteria.

<u>Surgical preparation</u>. The surgical preparation consisted of three main procedures – (1) head bar implantation, (2) AAV intracranial injection, and (3) GRIN lens implantation. In 6 animals (M0026, M0031, M0032, M0036, T01, T02), the preparation consisted of a single surgical event with all the procedures performed at once in the specified order. In the remaining 10 (M1027, M1035, M1039, M1050, M1051, M1052, M1053, M1058, M1062), the AAV injection was performed on an initial surgery, while the head bar and the GRIN lens implantation were performed on a second surgery typically 2-3 weeks later. No differences were observed between the two surgical protocols. For each surgical event, the animals were prepared as follows. One hour before the surgery mice were injected with Buprenorphine slow release (1 mg/Kg subcutaneous, 1 dose). The animals were later anesthetized using isoflurane (initially 2.5%, then 0.5-1.5%), injected with dexamethasone (2 mg/Kg intraperitoneal, 1 dose) and saline (0.3-0.4 mL, subcutaneous). Temperature was maintained by a heating pad, and breathing was checked regularly. After deep anesthesia was ensured, the animals were placed in the stereotaxic position, the skin was opened along the anterior-posterior axes and the periosteum was gently swabbed away. The skin was glued on the lateral sides, using Vetbond (3M), to leave enough bone expose for placing the head bar. To ensure a solid attachment of the head bar, a shallow grid was incised over the bone using a scalpel. Later, a custom made titanium headplate was secured on top of the skull using dental cement (C&B Metabond, Parkell).

<u>Viral injection</u>. The AAV injection was performed using a Hamilton Neuros 7000 series syringe (Hamilton Inc.). In every injection, we administered 0.5 µL of AAV-Syn-GCaMP6f mixed with an additional 0.5 µL of another liquid (i.e., the total injected volume was 1 µL, final dilution of 1:2; **Suppl. Table 2**). In some cases, the total volume (and dilution) was obtained adding 0.5 µL of Phosphate Buffered Saline (PBS); in other cases, we added a second virus to express tdTomato in a conditional dependent or retrograde way (**Suppl. Table 2**). In one case (M0026), we injected 0.5 µL of AAV-Syn-GCaMP6f-2A-tdTomato, mixed with 0.5 µL of PBS. To access the brain and perform the intracranial AAV injection the craniotomy was executed using trephine for microdrill (diameter = 1.8 mm, Fine Science Tools, #18004-18) and using standard procedures^52^. The craniotomy was centered on the target location (AP: 2.7 ± 0.1, ML: 1.35 ± 0.15 mm from Bregma, **Suppl. Table 1**). After the bone was removed, two injection sites were identified: each location was shifted of 0.2 ± 0.1 mm on both AP and medio-lateral (ML) axes and opposite directions. For example, if the target was AP: 2.7 and ML: 1.35 from Bregma, the first injection was performed in AP: 2.85, ML: 1.65 mm, the second in AP: 2.45, ML: 1.25 mm. Each location was further divided in 5 steps, spaced of 0.05 mm along the DV axis. For each step, 0.1 µL was injected at 20-40 nL/min and 3 minutes were waited before proceeding to the next step. To minimize tissue displacement, the most ventral site was always injected first, and the remaining sites followed serially. If the GRIN lens was implanted on a separate surgical event, the craniotomy was covered with the bone and/or Kwik-Sill (World Precision Instruments). Otherwise, the syringe was simply removed, and the GRIN lens was implanted.

<u>GRIN lens implantation</u>. The GRIN lens (ProView lens probe, 1 mm diameter x 4 mm length, #1050-004605, Inscopix Inc.) was initially cleaned. Then, it was attached to a dedicated holder from Inscopix and centered on the target location determined during the craniotomy using a stereotaxic arm. Before the insertion, a horizontal incision to the exposed craniotomy was performed. Typically, the scalpel was inserted 0.2-0.4 mm from the top of the tissue, hence penetrating both the dura and the cortex. This cut ensured the dura was opened, and prevented the tissue from bending and/or displacing during the initial phase of the insertion. If bleeding was observed, we used hemostatic sponge (Vetspon, Ferosan); once the bleeding stopped, the lens was lowered, with a speed of 0.1-0.2 mm/min to its final DV coordinate (corrected by ∼0.2 mm to compensate for the working distance of the lens, see **Suppl. Table 1**). When the GRIN lens was in its final position, it was cemented using opaque dental cement (C&B Metabond, Parkell). Around the lens, we later built a circular wall of dental cement (Stoelting) mixed with charcoal, to protect the lens from possible mechanical forces and from sidelights. The top of the lens was covered using Kwik-Cast (World Precision Instruments).

<u>Cre activation</u>. For the experiments in **Fig.3**, we expressed tdTomato in a Cre-dependent way. To do so, the two lines of animals required Cre activation that was performed between 6 to 8 weeks after the viral injection. For the strain Rasgrf2-2A-dCre (JAX 022864), the activation was performed using Trimethoprim (TMP). Specifically, TMP powder (T7883, Sigma) was dissolved in Dimethyl Sulfoxide (DMSO, D8418, Sigma) to a concentration of 100 mg/ml. Animals were injected intraperitoneally with the solution to a finale dosage of 300 mg/g of body weight on two different days, with one day in between the two injections^53^. The solution was made fresh every day before the injection. For the strain ER81-CreERT2 (JAX 013048), the Cre activation was achieved by the use of Tamoxifen. Tamoxifen (T5648, Sigma) was dissolved in corn oil (C8267, Sigma) at 55° C to a concentration of 30 mg/ml. Animals were injected subcutaneously with the solution, for a dosage of 180 mg/Kg, for five consecutive days. The tamoxifen/corn oil solution was made at the beginning of the injection cycle and stored in a dark and dry environment at 4°C for the remaining four days.

<u>Histology</u>. At the end of the imaging experiments, we verified the GRIN lens position by histology. The animals were deeply anesthetized with a cocktail of ketamine/xylazine, then perfused with PBS and 4% paraformaldehyde (PFA). The brain was extracted, placed in PFA overnight, then transferred to a solution of 30% sucrose in PBS, After the brain sunk, it was moved in an optimal cutting temperature compound (OCT) and frozen at −80° C. Subsequently, the brain was sliced (approximately 30 µm sections) with a low-temperature Cryostat (Leica Biosystems) and pasted on a glass, then stained with DAPI mounting solution (Vectashield, Vector Laboratories). Sections were examined and photographed under a fluorescence microscope (Leica DMI6000 B microscopy). The sample was imaged with an excitation wavelength of 358 nm for DAPI, 488 nm for GCaMP, and 554 nm for tdTomato. For anatomy experiments focused on L1 (**Suppl. Fig.9**), myelinated axons were visualized by staining in flat mounted cortex. Adult mice were perfused with PBS followed by 1% PFA. Cerebral cortex was separated from the rest of the brain, flat mounted, postfixed in 4% PFA, cryoprotected in 30% sucrose in PBS and sectioned on a freezing microtome in the tangential plane at 40 µm. Free-floating sections were stained with an antibody against myelin basic protein (monoclonal mouse-anti-myelin basic protein antibody, BioLegend #SMI-99P, 1:1000)). Myelin was visualized with a donkey-anti-mouse IgG, labeled with Cy5 (Jackson ImmunoResearch Labs #715-175-150, 1:500). Sections were mounted with ProLong Gold Antifade (Thermofisher) and imaged with an inverted microscope (Nikon, Eclipse Ti2) equipped with a CMOS camera at 10 x (Plan Apo 10x/0.45 objective) magnification.

### Choice task

<u>Trial structure</u>. The behavioral apparatus was custom-built and it was operated by custom-written code in the LabView environment. During the experiments, mice were head-fixed and restrained in a plastic tube. Two odor delivery ports and two liquid spouts were placed on the left and on the right of the animal head, and close to the mouth. In each session, the animal chose between two juices offered in variable quantities, delivered from left and right lick ports. Offers were represented by olfactory stimuli. Each juice flavor was associated with an odor, and the juice quantity was represented by the corresponding concentration. Before each trial, a vacuum sweep removed the odors remaining from the previous trial. Immediately thereafter, the two offers were presented simultaneously from two directions (left, right). Offer presentation lasted for 2.4-2.8 s (the time was fixed for each animal and varied across animals). After offer presentation, the animal indicated its choice by licking one of the two spouts. The liquid spouts were fixed and did not retract during odor presentation. The response period started with an auditory ‘go’ sound (0.2 s) and licking before the cue was disregarded. Licking was detected by two custom-made licking (touch) detectors, each connected with a single spout^54^. If the animal did not respond within 5 s, the trial was aborted and baited. Forced choice trials in which the animal licked the wrong spout were considered errors. Upon error trials, we presented the animals with white noise typically for 2-4 s (fixed for each session), we aborted the trial and repeated the same offer in the subsequent trial.

<u>Juice flavors and olfactory stimuli</u>. A juice pair was selected for every session from the following: (A) 15% sucrose vs. (B) water; (A) apple juice vs. (B) blueberry juice; (A) apple juice vs. (B) water; (A) 15% sucrose vs. (B) cranberry juice; (A) 15% sucrose vs. (B) grape juice; (A) water vs. (B) apple juice; (A) pomegranate juice vs. (B) water; (A) apple juice vs. (B) white grape juice; (A) white grape juice vs. (B) water; (A) grape juice vs. (B) cranberry juice; (A) 15% sucrose vs. (B) cranberry juice; (A) 15% sucrose vs. (B) peppermint tea; (A) elderberry juice vs. (B) white grape juice; (A) elderberry juice vs. (B) water; (A) 15% sucrose vs. (B) elderberry juice. In all sessions we used two odors, namely octanal (Sigma #0568) and octanol (Sigma #472328). We chose these odors because they are volatile, distinguishable^55^ and neutral (i.e., neither attractive nor aversive for mice^56^). The association between the juices and the odors varied pseudo-randomly across mice. The juice quantity offered to the animal varied between 0 and 7 drops, on 4 or 5 levels fixed for each session. The quantity was monotonically related to the represented odor concentration, which varied roughly linearly on a log2 scale (e.g., odor levels 1, 2, 4 and 8 ppm representing quantities 1, 2, 3 and 4 of juice). In each session, the left/right location of the two juices varied pseudo-randomly from trial to trial. Juice delivery took 150-1050 ms depending on the juice quantity (∼150 ms per quantum). Independent from the quantity, the reward period was kept constant within a session and was equal to the time necessary to deliver the maximum drops offered in that session (e.g., 900 ms for 6 drops, or 1050 ms for 7 drops). Odor presentation for the next trial started after a random period defined every trial from a uniform distribution typically varying between 3.8 and 4 s.

<u>Training</u>. As previously described^9^, animals were initially trained in a discrimination task; subsequently, we introduced the association between odor concentrations and reward quantities. When the animals reached ∼90% accuracy, we introduced the full task with binary choices. After full training, mice typically performed the task for 30-40 min each day, during which they completed 200-250 correct trials and received 0.8–1.6 ml of liquid reward. Sessions included ∼50% of forced choices (25% for each juice). To maintain the animal motivated for the experimental sessions, we alternated days with a session (or two, in some cases) as the one described, with days where with a session shorter and with higher number of forced choices.

### Analysis of choice data

All analyses of behavioral and neural data were conducted in Matlab (MathWorks). Behavioral choice patterns were analyzed using logistic regression, as previously described^14, 32^. First, we built a logistic model including the side bias, as follows:

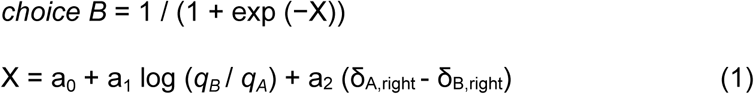

In Eq.1, *choice B* equals 1 if the animal chose juice B and 0 otherwise, *q_A_* and *q_B_* are the offered quantities for juices A and B, and δ_J,right_ = 1 if juice J was offered on the right and 0 otherwise (J = A or B). For each session, the logistic regression (maximum likelihood) returned two sigmoid functions with the same steepness and different flex points^32^. From fitted parameters a_0_, a_1_ and a_2_, we computed the choice accuracy *η* = a_1_, the (subjective) relative value of juices *ρ* = exp(−a_0_/a_1_), and the side bias *ε* = −a_2_/a_1_. Intuitively, *η* (also termed inverse temperature) was proportional to the sigmoid steepness and inversely related to choice variability, *ρ* was the quantity ratio *^q^_B_*^/*q*^*_A_* that made the animal indifferent between the two juices, and *ε* was a bias favoring one side over the other (specifically *ε* > 0 corresponded to a bias favoring the right side). Preliminary analyses^14^ indicated that the side bias was generally modest (|*ε*|<*ρ* in most sessions), but varied from session to session. Thus the analyses conducted here were restricted to sessions with a small side bias (see below), for which we ran a reduced logistic model:

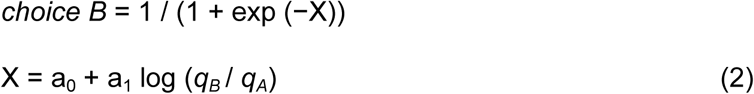

Again, from fitted parameters a_0_ and a_1_, we computed the choice accuracy *η* = a_1_ and the relative value *ρ* = exp(−a_0_/a_1_). As in previous studies^11, 14^, candidate variables possibly encoded by individual cells were defined based on the relative value measured in the same session.

### Two-photon calcium imaging

#### Imaging setup

Imaging was performed using an Ultima system (Prairie Technologies) built on an Olympus BX61W1 microscope. A tunable mode-locked laser (Mai Tai DeepSee Ti:Sapphire laser, Spectra-Physics) excited the tissue (wavelength = 920 nm for GCaMP6f, wavelength = 1040 nm for tdTomato). The fluorescence collected through a green filter (525/70 nm) and red filter (595/50), a GaAsp-PMT photo-detector with adjustable voltage gain, and offset feature (Hammatsu Photonics). The tissue was imaged with a long working distance 20x air objective with 0.45 NA, optimized for infrared wavelength transmission and with a correction collar (Olympus, AMEP4763EO).

#### Imaging procedures

Each field of view (FOV) was imaged with a resolution of 512×256 pixels and dwell time ≥0.3 µs. The region of interest (ROI) was drawn of different sizes for each sample to optimize the number of neurons and to obtain a minimum frame rate ≥7 Hz. The alignment between the behavioral system (in LabView) and the imaging system (Priare software, Bruker) was achieved as follows. For each trial, 2P scanning was triggered during the inter-trial interval (ITI), 2 s prior to the beginning of the odor delivery by a TTL signal from the behavioral system. A second trigger sent at the end of the reward delivery was used to stop the imaging sequence. Furthermore, to account for possible delays between the trigger TTL and the actual start of the imaging, another TTL from the imaging software (Priare software) was sent to the behavioral system (LabView) every trial and later used to align the data. From each animal we could typically image a volume of tissue that spanned 100-250 µm on the z axis (roughly corresponding to the DV axis). During the initial imaging session we performed Ca^2+^ imaging to qualitatively inspect the cortical volume. In this session we identified the experimental z planes (and in some cases portion of a plane) – i.e., the focal planes where we decided to perform imaging while the animal performed the choice task. These z planes are referred to as FOVs (3-12 FOVs per animal). Importantly, the FOVs were chosen blinded to their functional properties and such to avoid repetition of the same neurons across FOVs (minimum spacing of 20 µm in the z axis, if two FOVs were identified on the same Z plane an intermediate section wasn’t investigated). Typically, one FOV was imaged on each recording day.

#### Image processing

Acquired images in each session were analyzed independently. Raw imaging data were processed using CaImAn package in Matlab^33^. Images were first motion corrected on an average template of 500 frames. Then putative neurons were identified using Constrained Nonnegative Matrix Factorization (CNMF) to demix overlapping fluorescent signals and denoise the background^33, 57^. The quality of the segmentation was ensured by automatic controls implemented in CaImAn (threshold of the convolutional neural network, to evaluate the spatial footprint at 0.9; minimum signal to noise ratio, for the temporal profile of 1), and a manual inspection of the spatial and temporal components for each putative neuron (blind from their functional properties). The activity of each cell was then computed as ΔF/F, and we used this signal for all the other analysis (no spike deconvolution was performed).

#### Image registration

Most of the data included in this study are a subset of a longitudinal data set for which we imaged each FOV multiple time over the course of ≤20 weeks^14^. For each FOV and each pair of sessions, we registered images and matched neurons using a Bayesian procedure. First, we aligned the images of the two FOVs by projecting their neuronal footprints onto the same image and finding the rotations and translations that yield the highest cross-correlation between their projections. Then we modeled the distributions of centroid distances and spatial correlations as a weighted sum of the distributions of two subpopulations of cell-pairs, representing same cells and different cells. Using Bayes’ rule, we obtained the probability (Psame) for any pair of neighboring cells from different sessions to be the same cell, given their spatial correlation and centroid distance. Finally, we imposed a threshold of Psame>0.75. Cell pairs that satisfied this criterion were identified as matching. Additional details are provided elsewhere^14^.

### Classification of cells in L2/3 vs L5

#### Manual classification

The classification of neurons in “small” (L2/3) or “large” (L5) was performed manually, blind from the functional properties of neurons, and using morphological features. The experimenter inspected one FOV at the time, using both the time series and high spatial resolution images, to assign one of three possible labels: L2/3, L5, or not classifiable (NC). The same label was assigned to every neuron in the FOV. This procedure was done for two reasons. One, the most prominent feature of L5 cells was the apical dendrite. However, the dendrite is only present in pyramidal neurons. Assigning the same label to all the neurons helps mitigate the mislabeling of interneurons. Second, we typically chose FOVs with high morphological homogeneity, and no clear border between L2/3 and L5. In essence, we chose FOVs that we expected to be only (or mostly) in either L2/3 or L5. Last, FOVs with no clear border but high morphology heterogeneity were classified as NC. The accuracy was evaluated using two experimenters and comparing their results. The results were very consistent, and researchers agreed on 90% of the FOVs. In case the researchers didn’t agree, the FOV was labelled as NC. Of note, any mislabeling would most likely add noise and potentially hide differences across the two populations.

#### Automated classification

In four preparations, including one explicitly chosen to represent a hard case, we supplemented the manual classification with an automated analysis. We exploited the fact that segmentation yields a nonnegative factorization

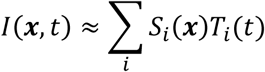

where *I*(***x***, *t*) is the raw image intensity at pixel location ***x*** (a 2-dimensional position) and frame *t*, *S*_*i*_ is the “spatial footprint” of the *ii*^th^ cell (in essence its average appearance) and is only a function of ***x***, and *T*_*i*_ is the temporal activity of the cell and is only a function of *tt*. The advantage of this representation is that it removes contamination from nearby cells, as each cell’s intensity is relegated to a separate component (*S*_*i*_, *T*_*i*_). For each cell’s spatial footprint *S*_*i*_, we identified the center-of-intensity (i.e., the intensity-weighted spatial average ***xx***) around which we snipped out a 19×19 region *S*^_*i*_(***x***). From the resulting intensity profile, we computed inner products with Zernike polynomials and used the resulting coefficients as a feature vector for classification. Specifically, if we define

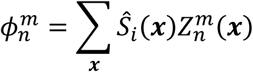

as the inner product with Zernike polynomial *Z*^*m*^_*n*_, then we defined

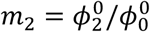

as the radial intensity cumulant (a measure of how intensity varies with radius) and

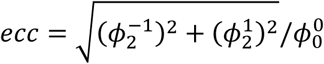

as the eccentricity of the intensity (zero for a perfectly circular cell). The Zernike polynomials were defined following the convention of Thibos et al^58^, and thus negative values of *m*_2_ in **Suppl. Fig.4** indicate that the intensity is concentrated towards the center of the ROI.

We found that the spatial footprints from CaImAn, given the settings we were using, did not accurately recapitulate cellular morphology as judged by a human observer. For this analysis, we recomputed a nonnegative matrix factorization from scratch using a custom pipeline for these four data sets. The algorithm will be published separately along with a larger package for the analysis of imaging data.

### Neuronal data set

The primary data set available for this study included 78 FOVs from 14 animals. This data set is nearly identical to a data set previously published^14^, except for two differences; (1) we included another animal (M0026); (2) we used more stringent behavioral criteria (see below).

Importantly, most FOVs were imaged multiple times over the course of many weeks (see above and [^14^]). We leveraged the richness of the whole data set in several ways. (1) We removed from the dataset all incomplete sessions – i.e., sessions that did not include forced choices for both juices, or such that the animal failed to choose each juice ≥80% of the time for at least some offer type. (2) We excluded from all analyses sessions with a large side bias (*ε*≥|1|). (3) The present study focuses on differences between L2/3 and L5. Most of our FOVs focused entirely on one layer, and only a fraction of FOVs included both L2/3 and L5. To avoid layer misclassifications, for the analysis of **Fig.4**, **Fig.6**, and **Fig.7**, we only included FOVs entirely focused on one layer (28 FOVs for L2/3; 27 FOVs for L5), for which imaging parameters had been suitably optimized. (4) As a first step in the neuronal analysis, we used procedures for variable selection to identify variables encoded in OFC (see below). For this analysis, we selected, for each FOV, the session with the largest number of neurons. (5) Building on the results of this analysis, we classified each cell as encoding one of 7 variables, namely *offer value A*, *offer value B*, *position of A*, *offer value I*, *offer value C*, *chosen value*, and *chosen side*. The classification was based on the total R^2^ summed across time windows that passed the ANOVA (see below). Previous work showed that this classification is rather noisy, because neuronal data are highly variable, the encoded variables are correlated, and the number of trials in any given session is limited^14^. At the same time, previous work showed that the representation of decision variables in OFC is longitudinally stable – i.e., neurons tend to encode the same variables in different sessions^14^. Thus to obtain a more robust classification, we used two sessions for each FOV (if available; see above, Image registration). Specifically, we chose the two sessions that maximized the number of matching neurons. The fraction of FOVs successfully registered was 24/28 for L2/3 and 22/27 for L5. In general, when a FOV was registered in 2 separate sessions, ∼50% of cells could be reliably identified in both sessions (matching neurons)^14^. For matching neurons, the classification was based on the total R^2^ summed across sessions and time windows that passed the ANOVA; for cells identified only in one session (non-matching neurons), the classification was based on the total R^2^ summed across significant time windows only for that session.

We also collected a smaller data set to specifically examine the effective connectivity between L2/3 and L5 using GCA (**Fig.5AB**; see below). This data set included 11 FOVs from 3 animals (M1051, M1053, and M1057; n = 2, 8, and 1 FOVs, respectively). The number of FOVs including both layers was relatively small because lenses were normally implanted to target preferentially one layer. That aside, a few considerations are in order. (1) To perform GCA across layers, L2/3 and L5 need to be imaged at the same time. Given the different brightness between the neurons of the two layers, PMT gain and laser power were tuned to allow the highest possible brightness that didn’t saturate L2/3 cells. As a result, L5 cells tended to be dimmer and potentially with a smaller dynamic range. (2) Each of these FOVs was imaged while the animal performed the task, but no behavioral threshold was imposed subsequently. (3) For the analysis, the data were initially processed as for the other dataset (see Image processing), but with one additional step. After the cell segmentation, in each FOV we manually labelled all L5 neurons. The labelling was based on the morphology and location within the FOV, and blind from the results of the GCA. All the remaining neurons were labelled as L2/3.

### Analysis of neuronal responses

#### Neuronal responses

. ΔF/F signals were examined in 5 time windows: *pre-offer* (600 ms preceding odor onset), *post-offer* (400-1000 ms from odor onset), *late delay* (1000-1600 ms from odor onset), *pre-choice* (600 ms preceding juice delivery onset), and *post-choice* (600 ms following juice delivery onset). An “offer type” was defined by two offered quantities. A “trial type” was defined by two offered quantities, their spatial configuration, and a choice. A “neuronal response” was defined as the activity of one neuron in one time window as a function of the trial type. Sessions typically included 10-12 offer types and 20-30 trial types. Neuronal responses were constructed by averaging the ΔF/F across trials for each trial type. Error trials in forced choices were excluded from the analysis. We also excluded from the analysis trial types with ≤2 trials.

#### Variable selection analysis

All the procedures for variable selection were as in previous studies^14^. Briefly, we conducted a series of analyses aimed to replicate previous results and to confirm the variables encoded in OFC. We proceeded in steps. First, the activity of each cell in each time window was examined by a 1-way ANOVA (factor: trial type). Neurons that passed the significance threshold (p<0.01) in ≥1 time window were identified as “task-related” and included in subsequent analyses. Second, we defined a series of variables that neurons in OFC might potentially encode (**Suppl. Table 4**). For each response passing the ANOVA criterion, we performed a linear regression against each variable. If the regression slope differed significantly from zero (p<0.05), the variable was said to “explain” the response. For each variable, the regression also provided an R^2^. For variables that did not explain the neuronal response, we arbitrarily set R^2^ = 0. Third, we constructed a population table. For each response passing the ANOVA criterion, we identified the variable that provided the best fit (highest R^2^). For each time window, we then computed the number of responses best explained by each variable. Fourth, we aimed to identify a small subset of variables that would best account for the whole neuronal population. To do so, we used a best-subset procedure, as in previous studies ^11, 59^. Importantly, this procedure is exhaustive and thus optimal. Responses from different time windows were pooled. For *n* = 1,2, …, all the possible subsets of *n* variables were considered, and the number of responses explained by each subset was computed. The best subset of *n* variables was that explaining the largest number of responses. The procedure was repeated for increasing *n*, until the best subset explained >80% of task-related responses. A variable analysis based on a stepwise procedure^11, 59^ provided identical results (not shown).

#### Cell classification

Confirming previous findings^14^, variable selection analyses indicated that neurons in OFC encoded variables *position of A*, *offer value A*, *offer value B*, *offer value I*, *offer value C*, *chosen side*, and *chosen value*. On this basis, we classified each neuron on the basis of the encoded variable. For each variable, each time window, and each session, we calculated the signed R^2^, where the sign was that of the regression slope. For each variable, we computed the sum(R^2^) as the total signed R^2^ across time windows. The encoded variable was that providing the highest ǁsum(R^2^)ǁ, where ǁ·ǁ indicates the absolute value. The sign of the encoding was that of sum(R^2^). Neurons that passed the ANOVA criterion but were not explained by any variable were classified as *untuned*. All subsequent analyses were based on this classification. For some analyses, we pooled *offer value A* cells and *offer value B* cells and collectively labeled them *offer value AB* cells. Similarly, we pooled *offer value I* and *offer value C* cells and collectively labeled them *offer value IC* cells.

### Analysis of spatial clustering

To assess whether neurons encoding any particular variable were spatially clustered within a cortical layer, we used an approach similar to that adopted in previous studies^60–62^. We first examined L2/3. We set a minimum distance *d_0_* = 20 μm and a search radius *d*. For each FOV and each cell, the position of the cell was computed as the center of mass of the 2D contour. We set a minimum distance *d_0_* and a search radius *d*. For each “reference” cell *i*, we examined neurons whose distance from *i* was larger than *d_0_* but smaller than *d* (see **Suppl. Fig.6**). We computed the number of such neurons encoding each of the 15 variables (7 signed variables + untuned), and we added these numbers to the table of cell counts (in the row corresponding to the variable encoded by cell *i*). We repeated this operation for each cell (i.e., each cell served as the reference cell) and for each FOV. Thus we obtained a table of cell counts for the whole population of L2/3. Of note, this table was symmetric because each pair of cells (*i*, *j*) contributed twice – once with *i* as the reference cell (row) and *j* as a cell in the vicinity of *i* (column), and once with *i* and *j* in reversed roles.

The table of cell counts was normalized by dividing each entry by the grand sum all the entries in the table. (Thus the sum of all the entries in the table of normalized cell counts equals 1.) We also computed the table of expected frequencies based on the fraction of cells encoding each variable across the entire population. In this table, entry (*u*, *v*) was equal to *p_uv_ = p_u_ p_v_*, where *p_x_* was the probability that any cell in L2/3 encoded variable *x*. (Thus the sum of all the entries in the table of expected frequencies equals 1.) Finally, we computed the disparity table defined as the entry-by-entry ratio (normalized cell count) / (expected frequency). In the disparity table, chance level =1; entries <1 or >1 indicated that cell counts were below or above chance level, respectively.

We repeated these operations varying *d* from 30 μm to 300 μm in 10 μm increments. To assess whether neurons encoding a particular variable formed spatial clusters, we examined how diagonal entries in the disparity table varied with *d*. Importantly, some of the variables were encoded by a small fraction of cells in L2/3. Consequently, the expected likelihood of finding two cells encoding one of these variables in the same FOV was close to zero. Thus the analysis of diagonal entries as a function of *d* was restricted to variables *u* for which the expected frequency was p*_uu_* ≥ 0.0002. We repeated this analysis for L5 setting *d_0_* = 30 μm and imposing the same criteria.

The minimum distance *d_0_* was imposed to address a concern due to possible errors during cell segmentation. In rare cases, a single neuron might be segmented as two separate cells, and such an occurrence would produce artifactual evidence supporting spatial clustering (because the two cells would likely encode the same variable). Considering that the typical cell size is 15-20 μm for L2/3 and 25-30 μm for L5, imposing a minimum distance *d_0_* = 20 μm for L2/3 and *d_0_* = 30 μm for L5 was considered safe. However, we also repeated the analysis imposing the more conservative criteria *d_0_* = 40 μm for L2/3 and *d_0_* = 60 μm for L5, and we obtained very similar results.

### Granger causality analysis

GCA was performed using the multivariate Granger causality toolbox (MVGC)^30^. For each session, time series data were the neuronal activity ΔF/F profiles aligned to the beginning of each trial and concatenated across trials. Time-domain Granger causality was calculated based on vector autoregressive modeling (VAR)^63–65^. To yield valid results, VAR coefficient have to be square, summable, and stable, and the MVGC toolbox provides suitable tests. All the sessions included in our analysis passed these criteria. The number of time lags (model order) was estimated independently for each session, as implemented in the package. Briefly, we fitted a VAR model to the entire session using different time lags (1 to 10), and then we selected the model providing the lowest Akaike Information Criterion (AIC). In our dataset, the model order ranged from 1 to 5 (71%,17%, 9%, 2%, and 1%, respectively). Ordinary least-squares was used to compute regression coefficients and the pairwise conditional causality was then calculated. For given source cell X and sink cell Y, the conditional causality was defined as the degree to which the past activity of X predicted the future activity of Y, over and above the prediction obtained from the past activity of Y and other conditional neurons. Conditional causality values that reached statistical significance (p < 0.01) after false discovery rate and multiple comparison correction were kept and others were set to zero. The output of the analysis was a binarized matrix for each session. The output was then sorted as needed.

#### Connections within and between layers

This analysis was restricted to FOVs including cells from L2/3 and L5 and presenting a clear border between layers (11 FOVs from 3 animals). For each FOV, we sorted cells in the GCA connectivity matrix so to obtain 4 sectors corresponding to connection patterns L2/3→L2/3, L2/3→L5, L5→L2/3, and L5→L5. For each sector, we computed the number of significant connections and the number of possible connections (the areas). In this calculation, we removed the diagonal because GCA does not provide auto-connections. For each sector, the population connection density was computed by summing all the significant connections across FOVs and dividing the result by the sum of all the areas across FOVs. To assess whether the connection density differed significantly from chance, we performed a bootstrap analysis. For each FOV, we kept fixed the areas and the number of significant connections, but we reassigned significant connections randomly to different pairs of cells. This procedure was repeated 10,000 times, and we obtained a distribution for the connection density expected by chance. If the measured connection density was below 2.5 or above the 97.5 percentile (p= 0.05, two tails) of this distribution, the departure from chance level was deemed statistically significant.

#### Connections within and between cell groups

A similar approach was used to examine connections within and between cell groups. For this analysis, however, we pooled FOVs from the two layers and we included FOVs that were not classified for a specific layer (78 FOVs from 14 animals). Whenever possible, cells were classified based on two sessions, as described above. For each FOV, we sorted cells in the GCA connectivity matrix based on the encoded variable (36 sectors). For each FOV and each sector, we computed the number of significant connections and the number of possible connections (excluding the diagonal). We combined the results at the population level and we computed the connection density for each sector. Finally, we ran a bootstrap analysis to assess whether, for each sector, the connection density differed significantly from chance (p<0.05, uncorrected, two tailed). To summarize the total connectivity of each cell group, we also computed the total connection density of the groups as source (column) and subtracted the total connection density of the groups as a sink (row).

### Comparing the representation of decision variables across cortical layers

To compare the representation of decision variables across layers, we used statistical analysis for categorical data^66^. Starting from a contingency table *X* whose entries were cell counts, we computed the corresponding table of odds ratios (ORs), as follows. For each location (i, j), we reduced the contingency table into a 2×2 matrix with elements

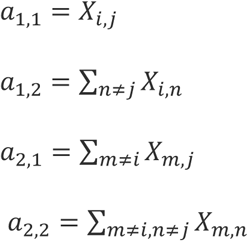

For each entry, the OR was defined as:

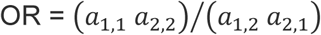

For each entry, OR = 1 was the chance level; conversely, OR > 1 and OR < 1 indicated that the cell count was larger or lower than expected by chance, respectively. Statistical significance from chance level was assessed using Fisher’s exact test (two-tailed)^66^.

### Activity profiles and strength of the encoding

To assess how of the representation of different decision variables evolved over time, we examined the activity profile of each cell group separately in the two layers. For this analysis, we pooled neurons encoding *offer value A* and *offer value B*, and neurons encoding *offer value I* and *offer value C*. However, cells with positive and negative encoding were examined separately (2 layers, 5 variables, 2 signs of encoding).

The analysis was based on the same data set used to compare the encoding of decision variables across layers (**Fig.6**). As described above, each cell was identified as encoding one of five variables, with a sign. If the classification resulted from multiple sessions (see above), neuronal activity recorded in each session contributed independently to the population average (i.e., most cells contributed to the average twice). Since the image acquisition rate could change from session to session, to compute the activity profiles, we interpolated Ca^2+^ traces to achieve a resolution of 1 ms. For each trial, neural signals (ΔF/F) were aligned to the offer onset and, separately, to the juice delivery. For neurons encoding a binary variable (*position of A*, *chosen side*), trials were divided in two groups according to the value of the encoded variable. For neurons encoding a non-binary variable (*offer value AB, offer value IC, chosen value*), trials were divided in three groups such that the encoded variable would be low, medium, or high (three bins of equal width). For each cell and each group of trials, we averaged ΔF/F across trials. Then we then computed the mean, standard deviation, and standard error across the population, for each value of the encoded variable. Thus we obtained the activity profiles shown in **Suppl. Fig.10** and **Suppl. Fig.11**.

Starting from the activity profiles, we also computed, the strength of the encoding (SOE). SOE was computed for each variable and for each sign of encoding, and was a function of time t in the trial. First, we computed the activity range (AR). For binary variables, AR was the difference between the two mean activity profiles. For non-binary variables, we regressed the 3 mean activity profiles against the encoded variable, AR was the activity range corresponding to the range of the encoded variable^67^. We also computed the standard deviation (SD) by averaging the standard deviations measured for the 2 or 3 values of the encoded variable, and the standard error SE = SD/sqrt(N), where N was the number of cells in the population. Finally, we defined SOE = AR/SE (**Suppl. Fig.12AB**).

For each variable, we also defined a combined SOE pooling neurons with the two signs of encoding, as follows: SOE = (sqrt(N^+^) AR^+^/SE^+^ + sqrt(N^–^) AR^–^/SE^–^) / sqrt(N^+^ + N^–^), where superscripts + and – indicate the sign of the encoding (**Fig.7GH**). To specifically assess differences in timing between layers and variables, we normalized each SOE trace by subtracting the value at offer onset and by dividing by baseline-subtracted trace by its maximum (**Fig.7I**).

## Acknowledgements

We thank Junxiao Hou for comments on the manuscript. This work was supported by the National Institutes of Health (grant numbers R21-DA042882 to C.P.S., R01-DA055709 to C.P.S., and R01-DC020034 to T.E.H.) and the McDonnell Center for System Neuroscience (Small Grant to A.L.).

## Contribution

A.L., M.Z., T.E.H. and C.P.S. designed the study; A.L., M.Z., M.C. and H.S. trained the animals; A.L and M.Z performed the experiments with assistance from M.C.; A.L. and M.Z. processed and analyzed the data; A.B. conducted anatomy experiments and helped with reconstruction and histology; C.P.S. and T.E.H. supervised the project; A.L. wrote the initial draft; all authors edited the manuscript;

Conflict of interest: Declared none

---

**Supplementary Figure 1.**
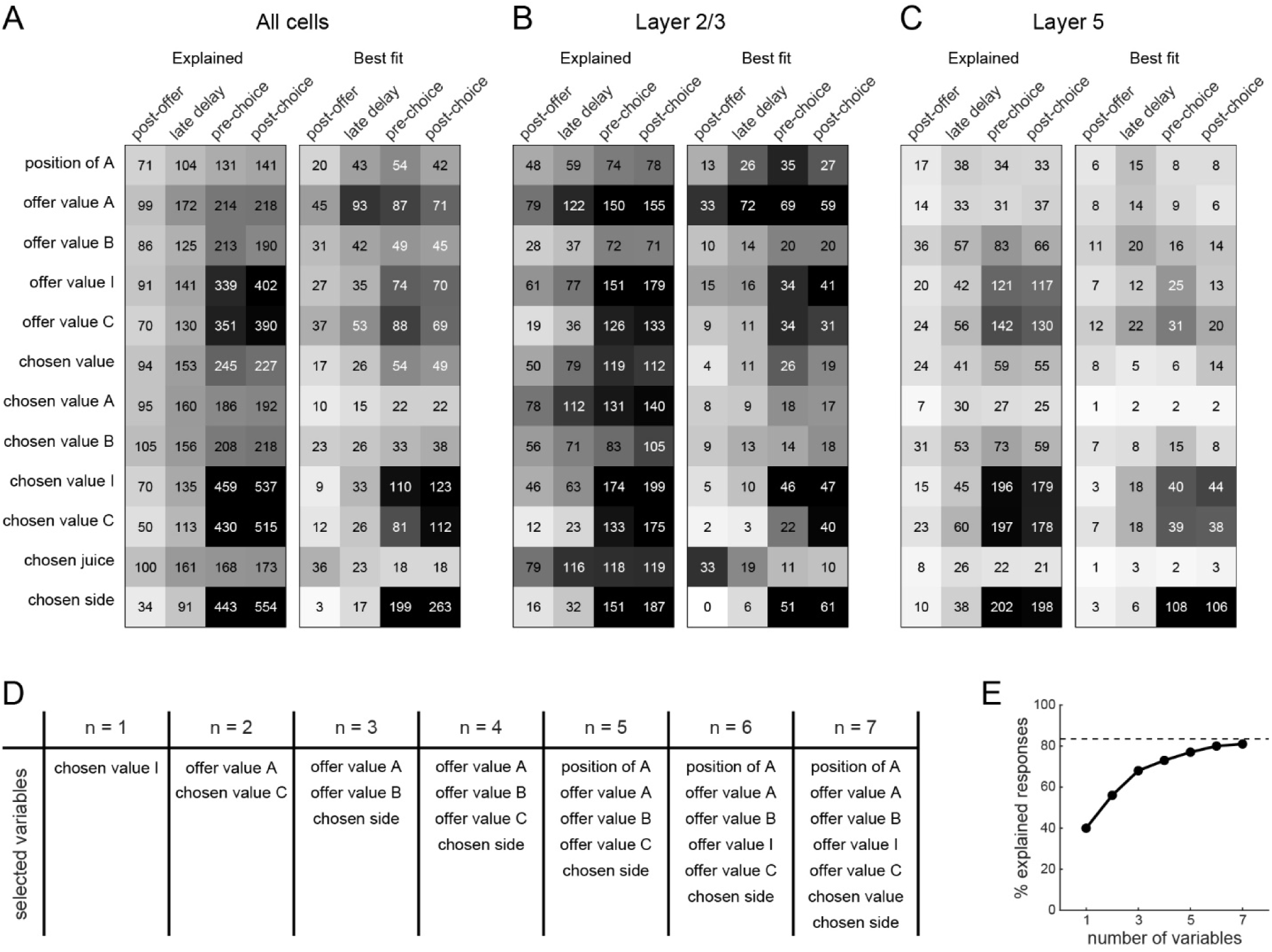
Encoding of decision variables in OFC. **A.** Population analysis, all cells. In both panels, rows and columns are candidate variables and time windows, respectively. In the left panel, entries are the number of responses explained by the variable in the corresponding time window. Gray shades recapitulate the cell counts. Of note, many responses are explained by ≥1 variables. In the right panel, entries are the number of responses for which the corresponding variable provides the best fit (highest R^2^). All neurons in the data set were included in this analysis (N = 7261; see **Methods**). **B.** Population analysis, layer 2/3 (N = 3134). **C.** Population analysis, layer 5 (N = 2027). **D.** Variable selection analysis, all cells. For n = 1…7, the table details the best subset of n variables as a function of n. **E.** Explained responses. The plot indicates the fraction of task-related responses explained by the best subset (y-axis) as a function of the number of variables included in the best subset (x-axis).

**Supplementary Figure 2.**
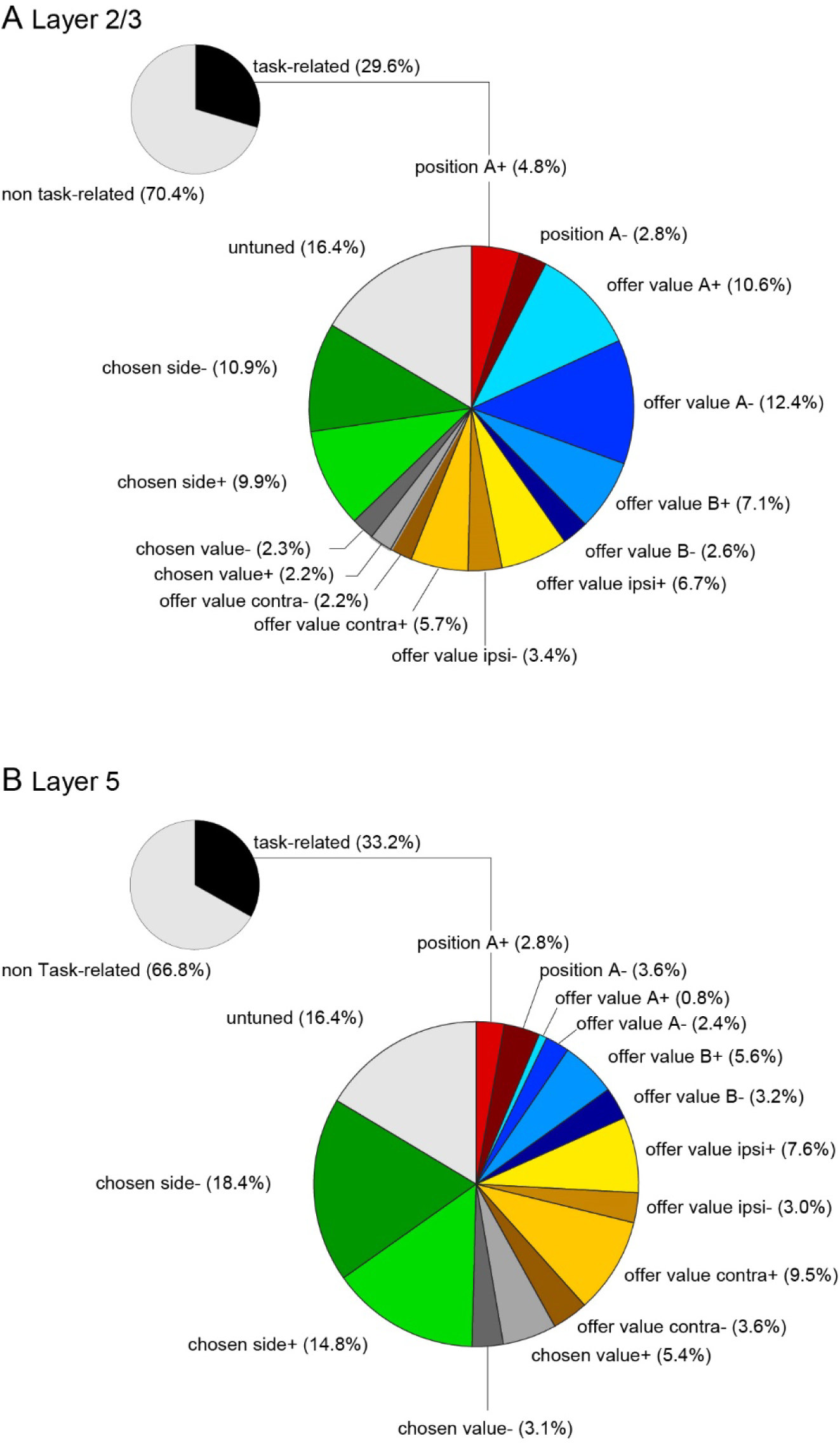
Fraction of neurons encoding different variables. **A.** All cells. The pie plot on the upper left shows the percentages of *task-related* cells and *non task-related* cells. The larger pie plot shows the percentages of *task-related* neurons encoding different variables. **B.** L2/3 cells. **C.** L5 cells. For each variable, we separated neurons with positive and negative encoding.

**Supplementary Figure 3.**
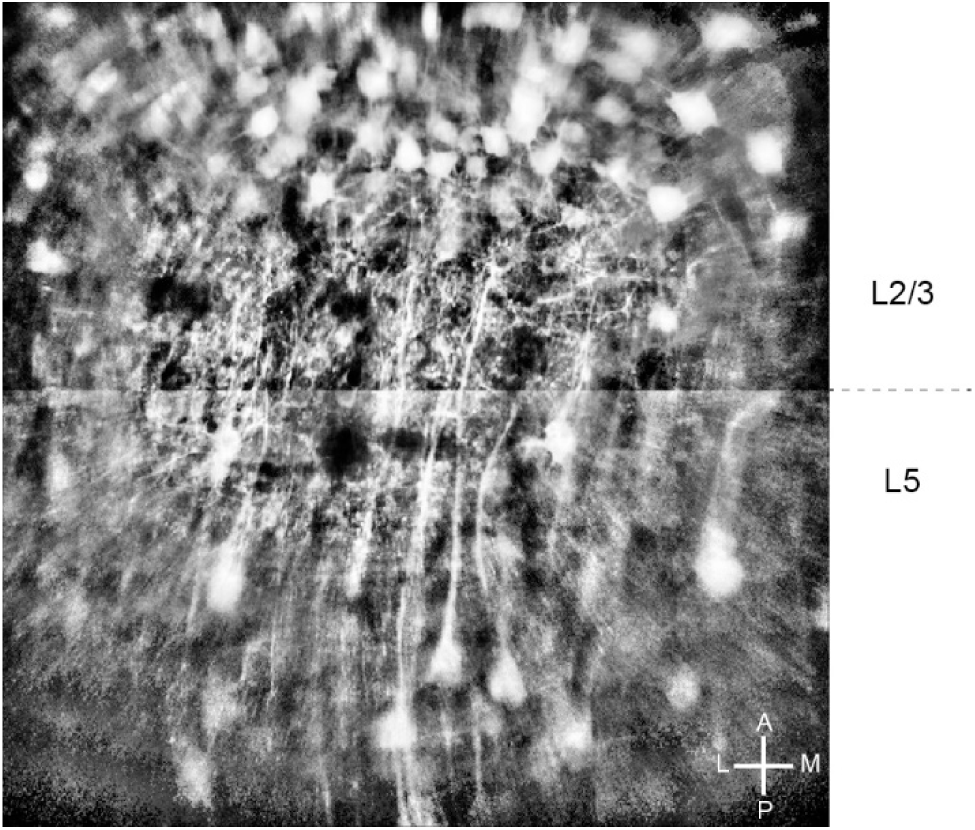
Example FOV, optimized visualization. Same example FOV illustrated in Fig.3AB, but here the contrast was independently and locally adjusted using CLAHE [^68^] for the lower and upper sections to better visualize the two clusters of neurons.

**Supplementary Figure 4.**
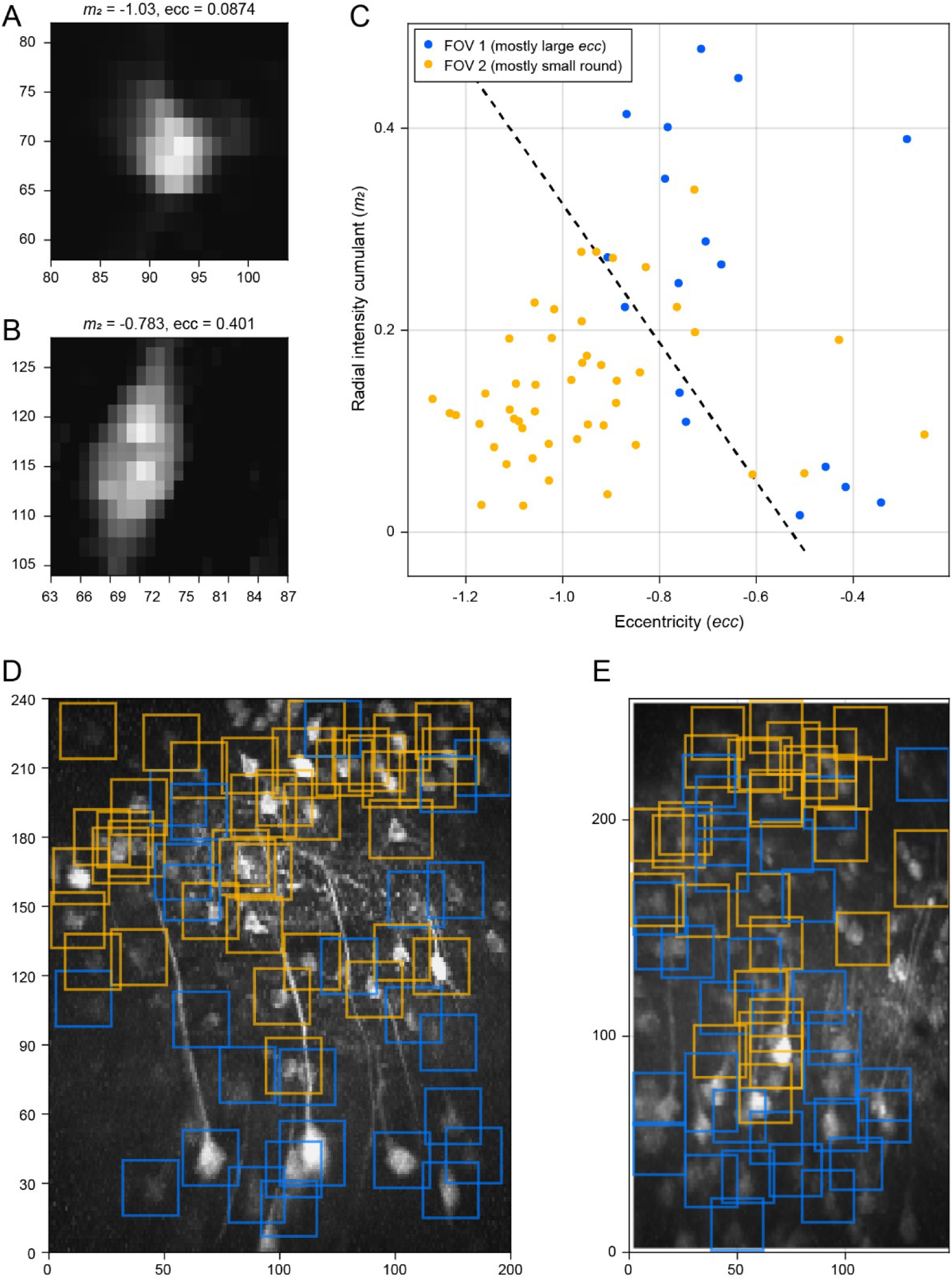
**Automated classification from cell morphology. AB**. 19 ×19 spatial footprints for neurons putatively from L2/3 (A) and L5 (B). Morphological parameters *mm*_2_ (radial intensity cumulant) and *ecc* (eccentricity) are indicated above each panel (see **Methods**). The L2/3 neuron is smaller (the intensity is concentrated more towards the center; i.e., *mm*_2_ is more negative) and round (eccentricity is small); the L5 neuron is larger (*mm*_2_ is larger) and highly eccentric. **C**. A scatter plot of *mm*_2_ and *ecc* for cells from two FOV used to train a classifier, each containing predominantly cells of only one layer. The color of each dot represents the FOV in which the cell was found. The dashed line represents a decision boundary computed from the optimal linear discriminant. **DE**. Classification of two held-out data sets (E was chosen deliberately as a hard case). Blue boxes (denoting putative L5 cells) tend to be towards the bottom, and yellow boxes (denoting putative L2/3 cells) tend to be towards the top, despite the classifier lacking spatial input.

**Supplementary Figure 5.**
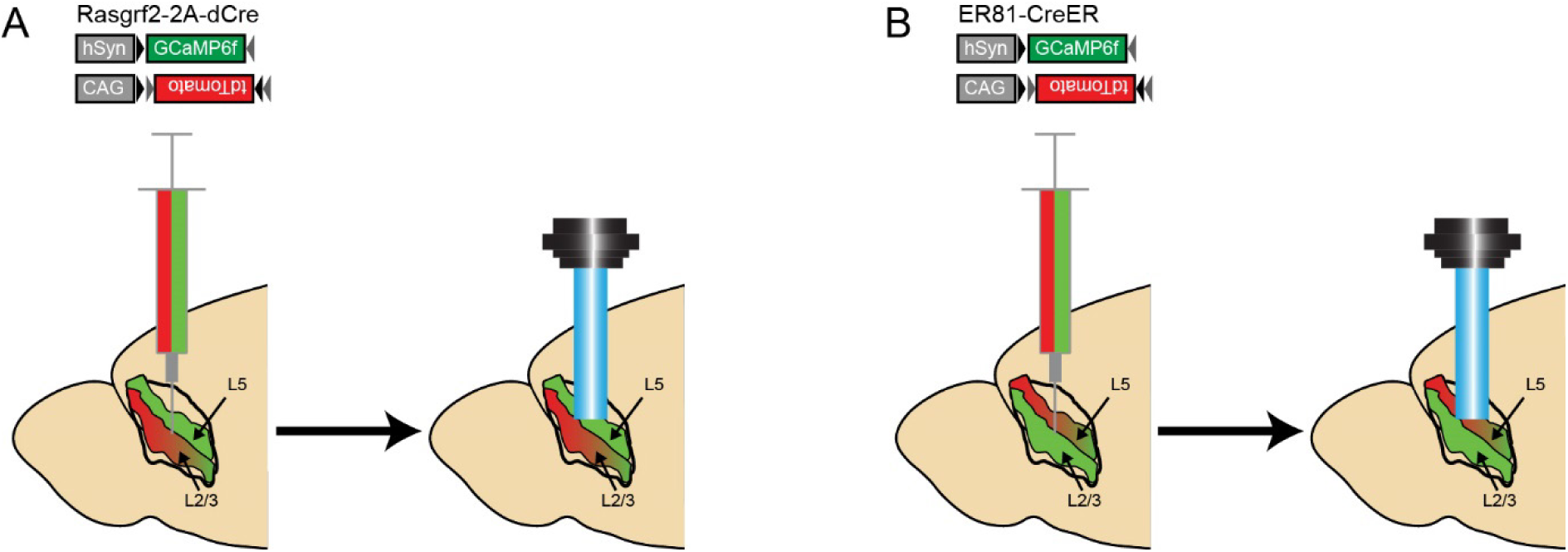
Layer-specific expression of red fluorescent protein tdTomato. The figure illustrates the procedure used to generate Fig.3. A mixture of two adeno-associate viruses (AAV) was injected intracranially in OFC. The viruses were used to express (1) GCaMP6f constitutively and (2) tdTomato in a Cre-dependent way (see **Methods**). After the injection, the GRIN lens was implanted in the infected cortical region. This dual-labeling procedure was performed in two different transgenic animals. **A.** Conditional expression of tdTomato in L2/3. We used Rasgrf2-2A-dCre mice that express inducible Cre (dCre) only in L2/3. dCre was then activated by injection of TMP. **B.** Conditional expression of tdTomato in L5. Here the experiment was conducted on ER-81CreER mice that express inducible Cre (CreER) only in L5. CreER was then activated by injection of tamoxifen.

**Supplementary Figure 6.**
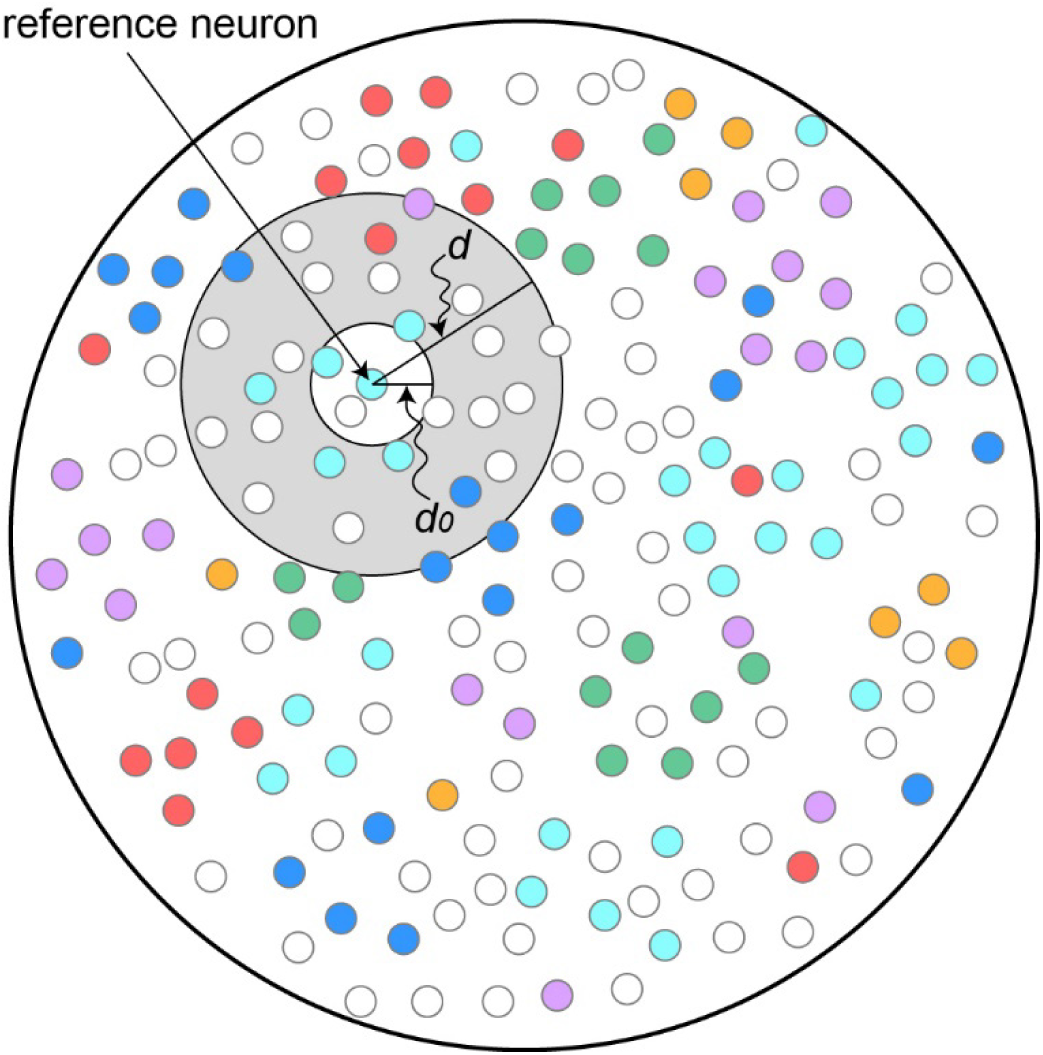
Procedure for the analysis of spatial clustering. The cartoon depicts a FOV. Cells encoding different variables are indicated with different colors. Each cell in the FOV served as reference neuron. For given reference neuron *i*, cells were said to be in the vicinity of *i* if their distance from *i* was between *d_0_* and *d*.

**Supplementary Figure 7.**
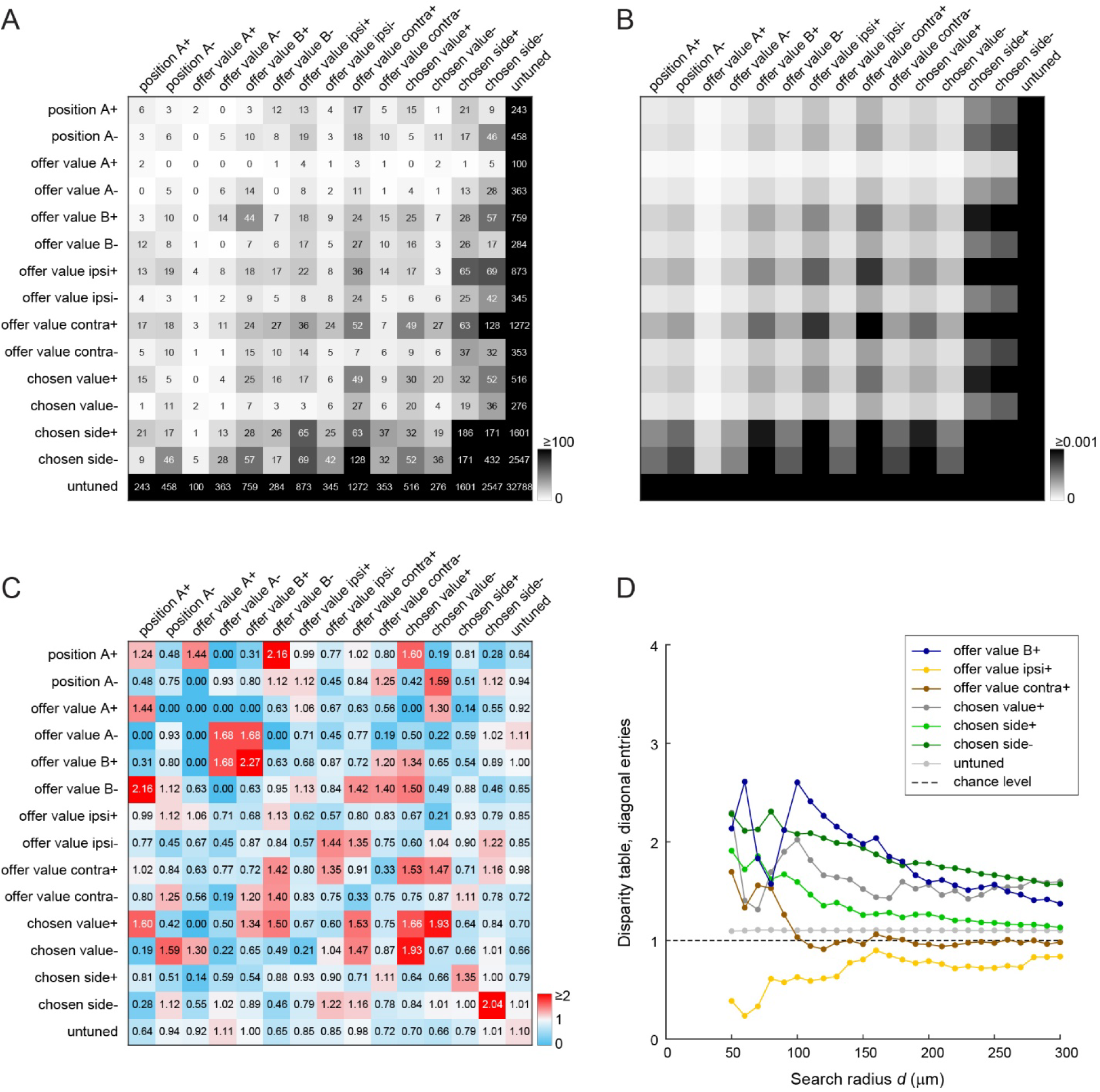
Spatial clustering of functional cell groups in L5. Same format as in Fig.4. In this case we set *d_0_* = 30 μm and *d* = 120 μm (see **Methods**). We computed cell counts (panel A), expected frequencies (panel B), and the disparity table (panel C). We then varied *d* between 50 μm and 300 μm (panel D). While the resulting picture was more noisy compared to that for L2/3 (because the neuronal population was smaller), most diagonal entries in the disparity table were >1 and decreased with distance.

**Supplementary Figure 8.**
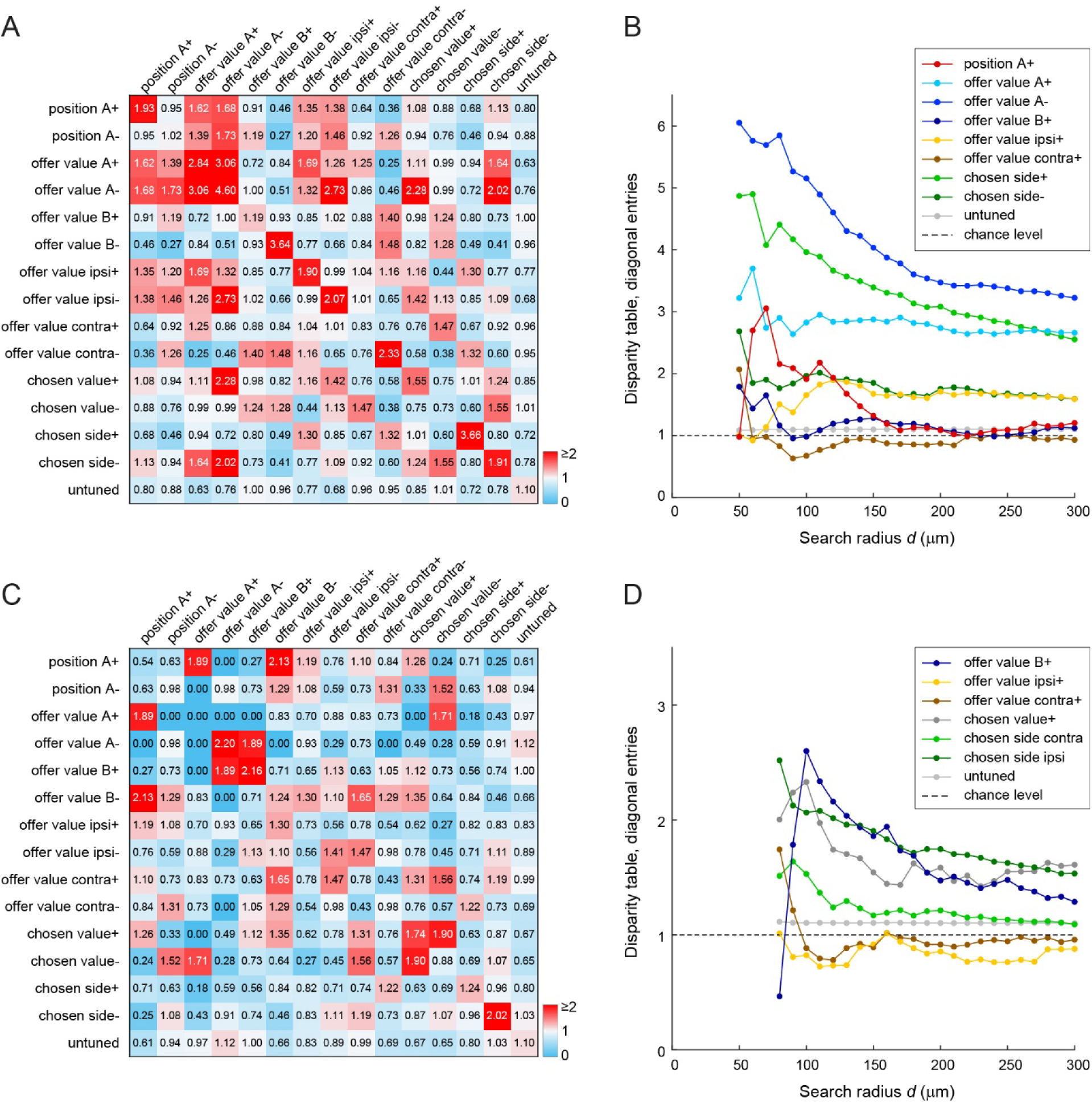
Spatial clustering, control analysis. The figure illustrates the results of the spatial clustering analysis obtained by setting *d_0_* = 40 μm for L2/3 (panels AB) and *d_0_* = 60 μm for L5 (panels CD).

**Supplementary Figure 9.**
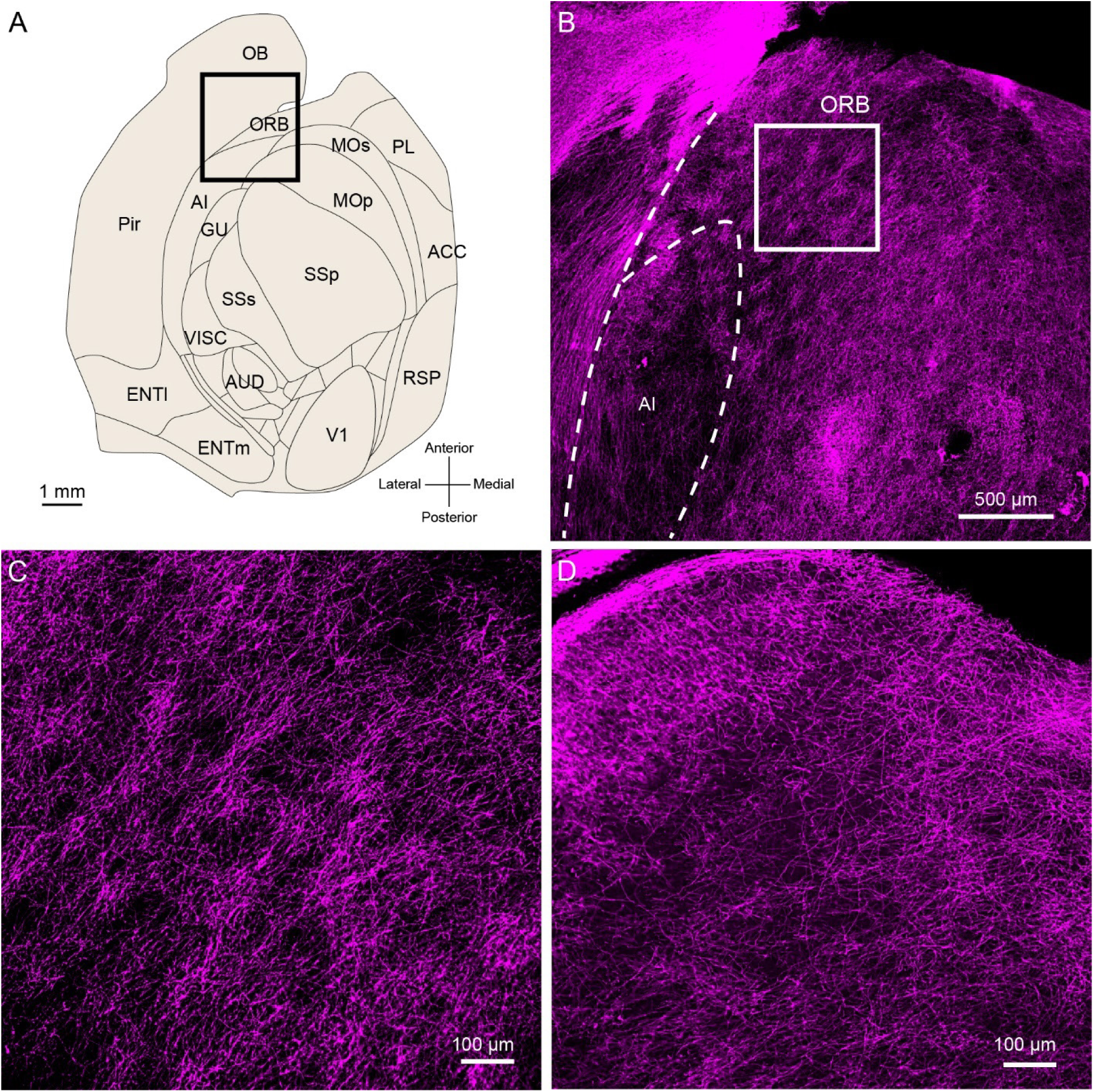
Patchy distribution of myelinated axons in L1 of orbitofrontal cortex. **A.** Area map. Map of areas in a flat-mounted left hemisphere of the mouse cerebral cortex. Abbreviations: anterior cingulate cortex (ACC), medial entorhinal cortex (ENTm), gustatory cortex (GU), primary motor cortex (MOp), secondary motor cortex (MOs), olfactory bulb (OB), piriform cortex (Pir), lateral prelimbic area (PL), posterior retrosplenial cortex (RSP), primary somatosensory area (SSp), secondary somatosensory area (SSs), primary visual area (V1), visceral cortex (VISC). **B.** Micrograph of boxed area outlined in panel A. The panel shows a non-uniform distribution of immuno-labeled myelinated axons in L1, 40 μm below the pial surface. **C.** Dense branching of myelinated axons in superficial L1. This panel illustrates the boxed area outlined in panel B. **D.** Sparsely branched myelinated axons in deep L1, 80 μm below the pial surface.

**Supplementary Figure 10.**
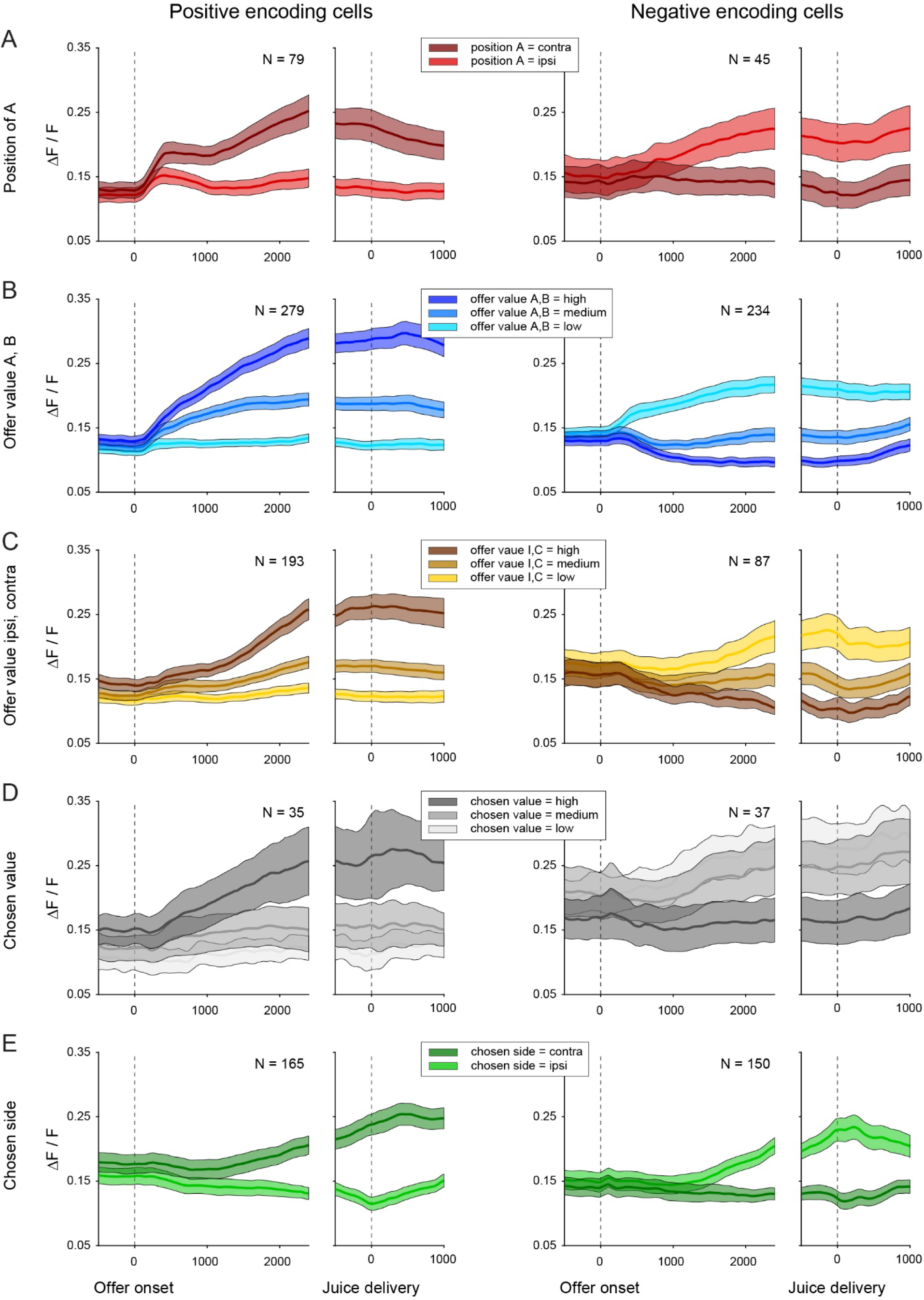
Activity profiles, layer 2/3. **A.** Cells encoding the variable *position of A*. We examined separately cells that presented higher activity for *position of A* = contra (positive encoding, left panels) and cells that presented higher activity for *position of A* = ipsi (negative encoding, right panels). For each cell, trials were divided depending on whether juice A was offered on the ipsi or contra side (*position of A* = ipsi or contra). Trials were aligned separately at offer onset and juice delivery. We examined neural signals (ΔF/F) and we obtained an activity profile for each neuron, and averaged the activity profiles across neurons (see **Methods**). Lines shown here are means and running error bars equal 2 s.e. for the relevant population. **B.** Cells encoding the variables *offer value AB*. We pooled neurons encoding variables *offer value A* and *offer value B*, but separated neurons with positive encoding (left panels) and negative encoding (right panels). For each cell, we divided trials in three groups depending on whether the relevant offer value was low, medium, or high (see **Methods**). We obtained an activity profile for each neuron and for each value bin, and we averaged the activity profiles across neurons for each value bin. Lines shown here are means and running error bars equal 2 s.e. for the population. **C.** Cells encoding the variable *offer value IC*. **D.** Cells encoding the variable *chosen value*. **E.** Cells encoding the variable *chosen side*. The number of cells included in each plot (N) is indicated in the corresponding panel. All times are in ms. All other conventions are as in panel A.

**Supplementary Figure 11.**
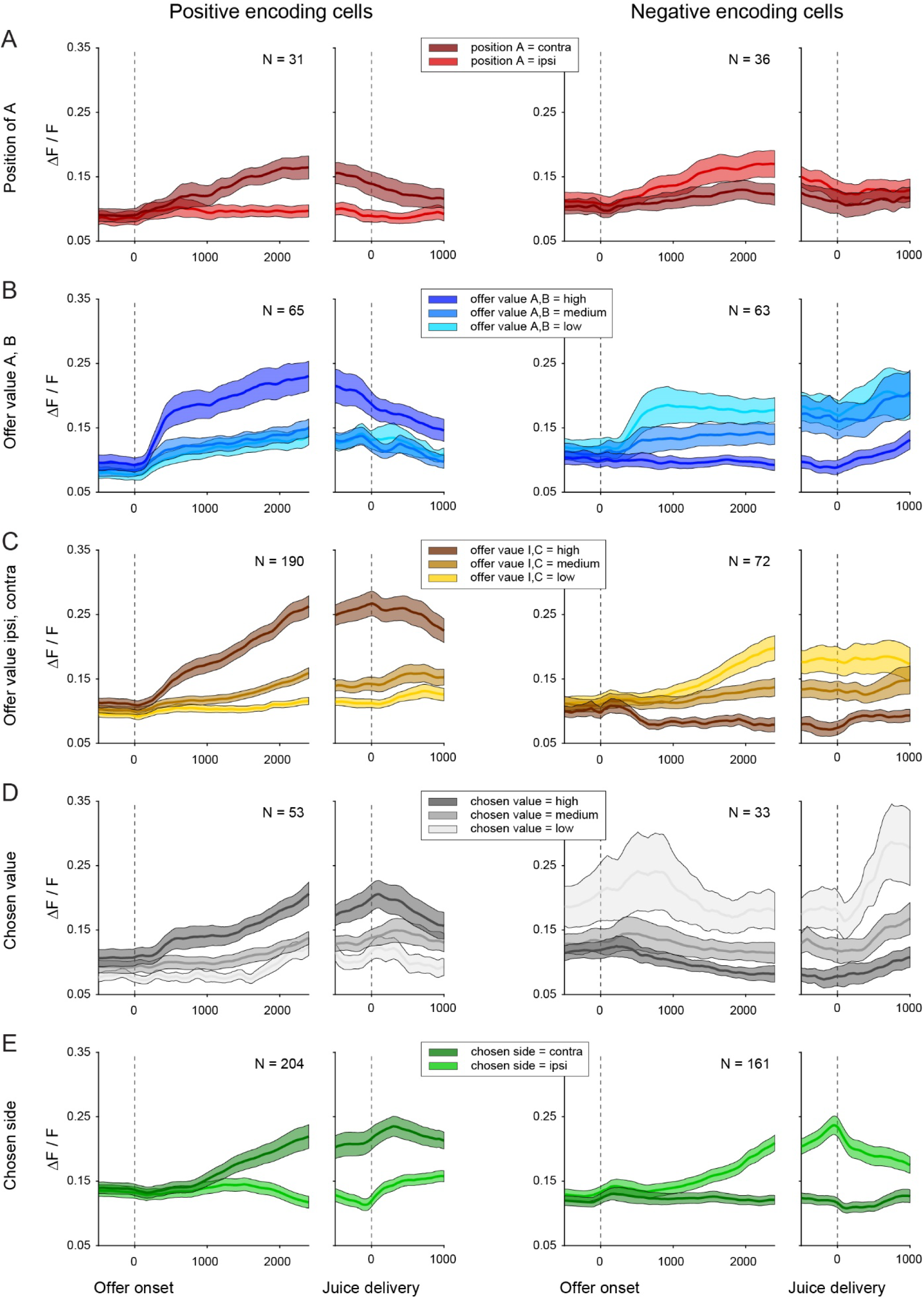
Activity profiles, layer 5. All conventions are as in **Suppl.** Fig.10.

**Supplementary Figure 12.**
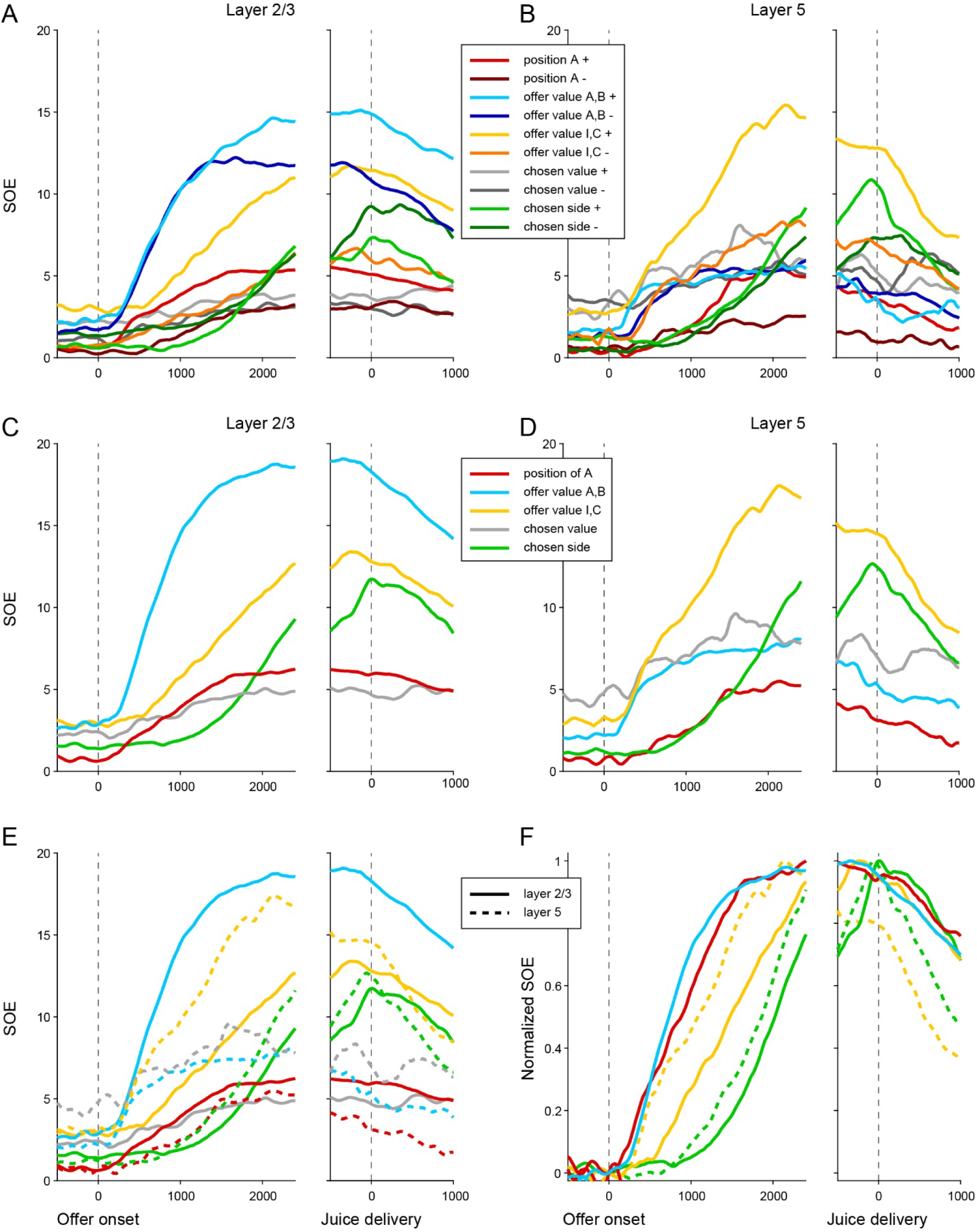
Strength of the encoding, detailed analysis. **A.** Layer 2/3, all variables. For each panel in **Suppl.** Fig.10, we computed the strength of the encoding (SOE). Traces shown here represent the SOE for each cell group (see color legend). **B.** Layer 5, all variables. Same analysis as in panel A for L5, based on **Suppl.** Fig.11. **CD.** Combined SOE. Traces shown here illustrate the combined SOE for each cell group (see color legend), separately for L2/3 (panel C) and L5 (panel D). **E.** Both layers. Traces in panels C and D are shown together (continuous lines = L2/3, dashed lines = L5). **F.** Both layers, normalized SOE. Traces in panel E vary both in amplitude and timing. To specifically assess differences in timing, we normalized each trace by subtracting the value at offer onset and by dividing by baseline-subtracted trace by its maximum. Panel F shows the normalized traces from the two layers for the most prominent variables. First to emerge after offer onset were variables *offer value AB* and *position of A* in L2/3. These were followed by variable *offer value IC* in L5 and, lastly, by variable *chosen side*. Note that both variables *offer value IC and chosen side* emerged in L5 first and were substantially delayed in L2/3 (by 200-300 ms), suggesting a feedback signal.

**Suppl. Table 1.** Animals included in the study. Target locations in OFC for each mouse used for neural recordings. Columns 4-6 (AP, ML, DV) indicate the distance from Bregma in mm. The last column (DVC) details the depth correction introduced to account for the working distance of the GRIN lenses, in mm. All lenses were placed in the left hemisphere. Animals A01, A02, and A03 were used for anatomy experiments focused on L1 (no GRIN lens implanted).

| Mouse ID | Strain | Sex | AP | ML | DV | DVC |
| --- | --- | --- | --- | --- | --- | --- |
| M0026 | C57BL/6J (JAX # 000664) | M | 2.6 | -1.5 | -3.1 | 0.15 |
| M0031 | Rasgrf2-2A-dCre (JAX # 022864) | M | 2.6 | -1.55 | -3.3 | 0.15 |
| M0032 | Rasgrf2-2A-dCre (JAX # 022864) | M | 2.6 | -1.55 | -3.3 | 0.15 |
| M0036 | PV-Cre (JAX # 008069) | M | 2.6 | -1.55 | -3.1 | 0.15 |
| M1027 | C57BL/6J (JAX # 000664) | F | 2.6 | -1.3 | -2.6 | 0.25 |
| M1035 | C57BL/6J (JAX # 000664) | F | 2.7 | -1.4 | -2.6 | 0.3 |
| M1039 | C57BL/6J (JAX # 000664) | F | 2.7 | -1.35 | -2.5 | 0.3 |
| M1050 | C57BL/6J (JAX # 000664) | M | 2.8 | -1.3 | -2.6 | 0.3 |
| M1051 | C57BL/6J (JAX # 000664) | F | 2.7 | -1.4 | -2.65 | 0.3 |
| M1052 | C57BL/6J (JAX # 000664) | F | 2.8 | -1.5 | -2.65 | 0.3 |
| M1053 | C57BL/6J (JAX # 000664) | M | 2.6 | -1.6 | -2.65 | 0.3 |
| M1057 | C57BL/6J (JAX # 000664) | F | 2.7 | -1.6 | -2.65 | 0.3 |
| M1058 | C57BL/6J (JAX # 000664) | F | 2.7 | -1.6 | -2.65 | 0.3 |
| M1062 | C57BL/6J (JAX # 000664) | M | 2.7 | -1.7 | -2.6 | 0.25 |
| T01 | ER81-CreERT2 (JAX # 013048) | F | 2.65 | -1.55 | -2.65 | 0.2 |
| T02 | Rasgrf2-2A-dCre (JAX # 022864) | M | 2.65 | -1.55 | -2.65 | 0.2 |
| A01 | C57BL/6J (JAX # 000664) | M |  |  |  |  |
| A02 | C57BL/6J (JAX # 000664) | F |  |  |  |  |
| A03 | C57BL/6J (JAX # 000664) | F |  |  |  |  |

**Suppl. Table 2.** Adeno-associated Viruses used in the experiments. Titers are expressed in GC/ml.

| **Virus** | **Addgene ID** | **Titer** |
| --- | --- | --- |
| pAAV.Syn.GCaMP6f.2A.tdTomato.WPRE.SV40 | #51085 | $2.5 \times 10^{13}$ |
| pAAV.Syn.GCaMP6f.WPRE.SV40 | #100837 | $2 \times 10^{13}$ |
| pAAV-FLEX-tdTomato | #28306 | $1.1 \times 10^{13}$ |
| pAAV.CAG.LSL.tdTomato | #100048 | $1.1 \times 10^{13}$ |
| Retrograde pAAVrg-CAG-tdTomato | #59462-r | $7 \times 10^{12}$ |

**Suppl. Table 3.** Results of ANOVA. N = 7261 cells were examined with a 1-way ANOVA (factor: trial type) and we imposed a significance threshold of p<0.01. Here rows are time windows and entries are the number of cells passing the statistical criterion. In total, 1804 cells (25%) passed the criterion in at least one time window. Neuronal responses that passed this test were identified as task-related and included in subsequent analyses. The right column of the table refers to the whole population. The table also includes separate cell counts for L2/3 (N = 3134) and L5 (N = 2027). Of note, some cells contributed to total cell count but were not included in either L2/3 or L5 (see **Methods**).

|  | All | L2/3 | L5 |
| --- | --- | --- | --- |
| pre-offer | 164 | 61 | 48 |
| post-offer | 368 | 183 | 103 |
| late delay | 559 | 267 | 185 |
| pre-choice | 1013 | 445 | 345 |
| post-choice | 1047 | 449 | 304 |
| at least 1 | 1804 | 763 | 581 |
| N cells total | 7261 | 3134 | 2027 |

**Suppl. Table 4.** Variables defined in the analysis of neuronal responses. In the analysis of neuronal data, we examined 12 variables. All values were expressed in units of juice B. The relative value *ρ* used to compute the unitary value of juice A was derived in each session from the logistic fit (**Eq.2**).

|  | Variable | Definition |
| --- | --- | --- |
| 1 | *position of A* | Binary; 0 if juice A was offered on the ipsilateral side, 1 otherwise |
| 2 | *offer value A* | Value of juice A offered = $\rho q_A$ |
| 3 | *offer value B* | Value of juice B offered = $q_B$ |
| 4 | *offer value I* | Value offered on the side ipsilateral to the recorded hemisphere |
| 5 | *offer value C* | Value offered on the side contralateral to the recorded hemisphere |
| 6 | *chosen value* | Value of the chosen juice |
| 7 | *chosen value A* | Chosen value if juice A was chosen, 0 otherwise |
| 8 | *chosen value B* | Chosen value if juice B was chosen, 0 otherwise |
| 9 | *chosen value I* | Chosen value if ipsilateral side was chosen, 0 otherwise |
| 10 | *chosen value C* | Chosen value if contralateral was chosen, 0 otherwise |
| 11 | *chosen juice* | Binary; 0 if juice A was chosen, 1 if juice B was chosen |
| 12 | *chosen side* | Binary; 0 if ipsilateral side was chosen, 1 otherwise |

**Supplementary Table 5.** Post-hoc analysis. We tested the selected variables (variable X) against highly correlated and thus competitive alternative variables (variable Y). Each row focuses on one pairwise comparison. We considered the enlarged subset including the three selected variables and variable Y. The marginal explanatory power of variable X, indicated with nX, was defined as the number of responses explained by the enlarged subset minus the number of responses explained when variable X was removed from the enlarged subset. Similarly, nY was the number of responses explained by the enlarged subset minus the number of responses explained when variable Y was removed from the enlarged subset. By definition of best subset, nX ≥ nY. To assess whether this inequality was statistically significant, we conducted a binomial test.

| Variable X | Variable Y | nX | nY | P value |
| --- | --- | --- | --- | --- |
| *offer value A* | *chosen value A* | 75 | 36 | $<10^{-4}$ |
| *offer value B* | *chosen value B* | 63 | 41 | 0.012 |
| *offer value ipsi* | *chosen value ipsi* | 61 | 29 | $<10^{-4}$ |
| *offer value contra* | *chosen value contra* | 88 | 18 | 0 |
| *chosen side* | *chosen value ipsi* | 84 | 29 | $<10^{-7}$ |
| *chosen side* | *chosen value contra* | 112 | 18 | 0 |

## Notes

### Competing Interest Statement

The authors have declared no competing interest.

## References

1. Padoa-Schioppa, C. & Conen, K.E. Orbitofrontal cortex: A neural circuit for economic decisions. Neuron 96, 736–754 (2017).

2. Rich, E.L., Stoll, F.M. & Rudebeck, P.H. Linking dynamic patterns of neural activity in orbitofrontal cortex with decision making. Curr Opin Neurobiol 49, 24–32 (2018).

3. Knudsen, E.B. & Wallis, J.D. Taking stock of value in the orbitofrontal cortex. Nat Rev Neurosci 23, 428–438 (2022).

4. Ballesta, S., Shi, W., Conen, K.E. & Padoa-Schioppa, C. Values encoded in orbitofrontal cortex are causally related to economic choices. Nature 588, 450–453 (2020).

5. Ballesta, S., Shi, W. & Padoa-Schioppa, C. Orbitofrontal cortex contributes to the comparison of values underlying economic choices. Nat Commun 13, 4405 (2022).

6. Camille, N., Griffiths, C.A., Vo, K., Fellows, L.K. & Kable, J.W. Ventromedial frontal lobe damage disrupts value maximization in humans. J Neurosci 31, 7527–7532 (2011).

7. Yu, L.Q., Dana, J. & Kable, J.W. Individuals with ventromedial frontal damage display unstable but transitive preferences during decision making. Nat Commun 13, 4758 (2022).

8. Lak, A., et al. Orbitofrontal cortex is required for optimal waiting based on decision confidence. Neuron 84, 190–201 (2014).

9. Kuwabara, M., Kang, N., Holy, T.E. & Padoa-Schioppa, C. Neural mechanisms of economic choices in mice. eLife 9 (2020).

10. Gore, F., et al. Orbitofrontal cortex control of striatum leads economic decision-making. Nat Neurosci 26, 1566–1574 (2023).

11. Padoa-Schioppa, C. & Assad, J.A. Neurons in orbitofrontal cortex encode economic value. Nature 441, 223–226 (2006).

12. Padoa-Schioppa, C. Neuronal origins of choice variability in economic decisions. Neuron 80, 1322–1336 (2013).

13. Pastor-Bernier, A., Stasiak, A. & Schultz, W. Orbitofrontal signals for two-component choice options comply with indifference curves of revealed preference theory. Nat Commun 10, 4885 (2019).

14. 14. Zhang, M., et al. The representation of decision variables in orbitofrontal cortex is longitudinally stable. *Cell Reports* 43 (2024).

15. Rich, E.L. & Wallis, J.D. Decoding subjective decisions from orbitofrontal cortex. Nat Neurosci 19, 973–980 (2016).

16. Shi, W., Ballesta, S. & Padoa-Schioppa, C. Neuronal origins of reduced accuracy and biases in economic choices under sequential offers. Elife 11 (2022).

17. Balewski, Z.Z., Elston, T.W., Knudsen, E.B. & Wallis, J.D. Value dynamics affect choice preparation during decision-making. Nat Neurosci 26, 1575–1583 (2023).

18. Solway, A. & Botvinick, M.M. Goal-directed decision making as probabilistic inference: a computational framework and potential neural correlates. Psychological Review 119, 120–154 (2012).

19. Rustichini, A. & Padoa-Schioppa, C. A neuro-computational model of economic decisions. J Neurophysiol 114, 1382–1398 (2015).

20. Friedrich, J. & Lengyel, M. Goal-directed decision making with spiking neurons. J Neurosci 36, 1529–1546 (2016).

21. 21. Song, H.F., Yang, G.R. & Wang, X.J. Reward-based training of recurrent neural networks for cognitive and value-based tasks. in *Elife* (2017).

22. Zhang, Z., Cheng, Z., Lin, Z., Nie, C. & Yang, T. A neural network model for the orbitofrontal cortex and task space acquisition during reinforcement learning. PLoS Comput Biol 14, e1005925 (2018).

23. Battista, A., Padoa-Schioppa, C. & Wang, X.J. Neural Circuit Underlying Economic Decisions: Insights from a Computational Model. in *Society for Neuroscience Annual Meeting* (Chicago (IL), 2024).

24. Douglas, R.J. & Martin, K.A. Neuronal circuits of the neocortex. Annu Rev Neurosci 27, 419–451 (2004).

25. Harris, K.D. & Shepherd, G.M. The neocortical circuit: themes and variations. Nat Neurosci 18, 170–181 (2015).

26. Douglas, R.J. & Martin, K.A. Recurrent neuronal circuits in the neocortex. Curr Biol 17, R496–500 (2007).

27. Ko, H., et al. Functional specificity of local synaptic connections in neocortical networks. Nature 473, 87–91 (2011).

28. Cossell, L., et al. Functional organization of excitatory synaptic strength in primary visual cortex. Nature 518, 399–403 (2015).

29. Bressler, S.L. & Seth, A.K. Wiener-Granger causality: a well established methodology. Neuroimage 58, 323–329 (2011).

30. Barnett, L. & Seth, A.K. The MVGC multivariate Granger causality toolbox: a new approach to Granger-causal inference. J Neurosci Methods 223, 50–68 (2014).

31. Seth, A.K., Barrett, A.B. & Barnett, L. Granger causality analysis in neuroscience and neuroimaging. J Neurosci 35, 3293–3297 (2015).

32. Padoa-Schioppa, C. Logistic analysis of choice data: A primer. Neuron 110, 1615–1630 (2022).

33. Giovannucci, A., et al. CaImAn an open source tool for scalable calcium imaging data analysis. Elife 8 (2019).

34. Makino, H., et al. Transformation of cortex-wide emergent properties during motor learning. Neuron 94, 880–890 e888 (2017).

35. Kondo, M. & Matsuzaki, M. Neuronal representations of reward-predicting cues and outcome history with movement in the frontal cortex. Cell Rep 34, 108704 (2021).

36. Sheikhattar, A., et al. Extracting neuronal functional network dynamics via adaptive Granger causality analysis. Proc Natl Acad Sci U S A 115, E3869–E3878 (2018).

37. Gilbert, C.D. & Wiesel, T.N. Columnar specificity of intrinsic horizontal and corticocortical connections in cat visual cortex. J Neurosci 9, 2432–2442 (1989).

38. Rossi, L.F., Harris, K.D. & Carandini, M. Spatial connectivity matches direction selectivity in visual cortex. Nature 588, 648–652 (2020).

39. Jordan, R. & Keller, G.B. Opposing influence of top-down and bottom-up input on excitatory layer 2/3 neurons in mouse primary visual cortex. Neuron 108, 1194–1206 e1195 (2020).

40. Olsen, S.R., Bortone, D.S., Adesnik, H. & Scanziani, M. Gain control by layer six in cortical circuits of vision. Nature 483, 47–52 (2012).

41. Adesnik, H. & Scanziani, M. Lateral competition for cortical space by layer-specific horizontal circuits. Nature 464, 1155–1160 (2010).

42. Pluta, S.R., Telian, G.I., Naka, A. & Adesnik, H. Superficial layers suppress the deep layers to fine-tune cortical coding. J Neurosci 39, 2052–2064 (2019).

43. Anderson, C.T., Sheets, P.L., Kiritani, T. & Shepherd, G.M. Sublayer-specific microcircuits of corticospinal and corticostriatal neurons in motor cortex. Nat Neurosci 13, 739–744 (2010).

44. Masamizu, Y., et al. Two distinct layer-specific dynamics of cortical ensembles during learning of a motor task. Nat Neurosci 17, 987–994 (2014).

45. Finn, E.S., Huber, L., Jangraw, D.C., Molfese, P.J. & Bandettini, P.A. Layer-dependent activity in human prefrontal cortex during working memory. Nat Neurosci 22, 1687–1695 (2019).

46. Conen, K.E. & Padoa-Schioppa, C. Neuronal variability in orbitofrontal cortex during economic decisions. J Neurophysiol 114, 1367–1381 (2015).

47. Daie, K., Svoboda, K. & Druckmann, S. Targeted photostimulation uncovers circuit motifs supporting short-term memory. Nat Neurosci 24, 259–265 (2021).

48. Onken, A., Xie, J., Panzeri, S. & Padoa-Schioppa, C. Categorical encoding of decision variables in orbitofrontal cortex. PLoS Comput Biol 15, e1006667 (2019).

49. Hirokawa, J., Vaughan, A., Masset, P., Ott, T. & Kepecs, A. Frontal cortex neuron types categorically encode single decision variables. Nature 576, 446–451 (2019).

50. Cai, X. & Padoa-Schioppa, C. Contributions of orbitofrontal and lateral prefrontal cortices to economic choice and the good-to-action transformation. Neuron 81, 1140–1151 (2014).

51. Guo, Z.V., et al. Procedures for behavioral experiments in head-fixed mice. PLoS One 9, e88678 (2014).

52. Resendez, S.L., et al. Visualization of cortical, subcortical and deep brain neural circuit dynamics during naturalistic mammalian behavior with head-mounted microscopes and chronically implanted lenses. Nat Protoc 11, 566–597 (2016).

53. Suzuki, M., Aru, J. & Larkum, M.E. Double-μPeriscope, a tool for multilayer optical recordings, optogenetic stimulations or both. Elife 10 (2021).

54. Slotnick, B. A simple 2-transistor touch or lick detector circuit. J Exp Anal Behav 91, 253–255 (2009).

55. Laska, M., Rosandher, A. & Hommen, S. Olfactory discrimination of aliphatic odorants at 1 ppm: too easy for CD-1 mice to show odor structure-activity relationships? J Comp Physiol A Neuroethol Sens Neural Behav Physiol 194, 971–980 (2008).

56. Saraiva, L.R., et al. Combinatorial effects of odorants on mouse behavior. Proc Natl Acad Sci U S A 113, E3300–3306 (2016).

57. Pnevmatikakis, E.A., et al. Simultaneous denoising, deconvolution, and demixing of calcium imaging data. Neuron 89, 285–299 (2016).

58. Thibos, L.N., et al. Standards for reporting the optical aberrations of eyes. J Refract Surg 18, S652–660 (2002).

59. Glantz, S.A. & Slinker, B.K. *Primer of applied regression & analysis of variance* (McGraw-Hill, Medical Pub. Division, New York, 2001).

60. Paik, S.B. & Ringach, D.L. Retinal origin of orientation maps in visual cortex. Nat Neurosci 14, 919–925 (2011).

61. Rodieck, R.W. The density recovery profile: a method for the analysis of points in the plane applicable to retinal studies. Vis Neurosci 6, 95–111 (1991).

62. Shin, J.H., Song, M., Paik, S.B. & Jung, M.W. Spatial organization of functional clusters representing reward and movement information in the striatal direct and indirect pathways. Proc Natl Acad Sci U S A 117, 27004–27015 (2020).

63. Rozanov, I.A. *Stationary random processes* (Holden-Day, San Francisco,, 1967).

64. Hamilton, J.D. Time series analysis (Princeton University Press, Princeton, N.J., 1994).

65. Lütkepohl, H. *New introduction to multiple time series analysis* (Springer, Berlin, 2005).

66. Agresti, A. An introduction to categorical data analysis (Wiley, Hoboken, NJ, 2019).

67. Padoa-Schioppa, C. Range-adapting representation of economic value in the orbitofrontal cortex. J Neurosci 29, 14004–14014 (2009).

68. Pizer, S.M., et al. Adaptive histogram equalization and its variations. Computer Vision, Graphics, and Image Processing 39, 355–368 (1987).

